# Dual-BONCAT Reveals Distinct Subpopulations of Anabolically Active Cells

**DOI:** 10.1101/2024.11.23.624985

**Authors:** Dylan Mankel, Yasheng Maierhaba, Claire Momjian, Federica Calabrese, Solange Duhamel, Jeffrey Marlow

## Abstract

Bio-Orthogonal Non-Canonical Amino Acid Tagging (BONCAT) has emerged as a prominent molecular technique that enables microbial ecologists to visualize and identify metabolically active cells in cultures and complex microbial communities. To date, researchers have used just one non-canonical amino acid (ncAA) in a given experiment; here, we validate a novel approach using two different ncAAs in a single experiment. This advancement facilitates the detection of differentially active subpopulations within the same experimental context, thereby reducing the uncertainty and variability associated with parallel treatments, and providing precise spatial information about organisms that are active under distinct conditions or at different times. We show that both ncAAs can be taken up by *E. coli* cultures and by constituents of the Little Sippewissett Salt Marsh microbiome, resulting in fluorescence signals that are significantly higher than background and ncAA-free control experiments, as well as differential labeling patterns reflective of distinct subpopulations. As a proof of concept, we implemented this “dual-BONCAT” approach in salt marsh sediments, adding one ncAA during daytime hours and the other at night. Subpopulations of cells that were anabolically active during the day and/or night were distinguishable by both fluorescence microscopy, and by fluorescence-activated cell sorting. Subsequent high-throughput 16S rRNA gene amplicon sequencing of active subpopulations revealed that *Methylobacterium*, potentially feeding on plant exudate carbon, was preferentially active during the day, while sulfur-cycling taxa dominated the night-active population. Dual-BONCAT offers an important advancement in multiplexing substrate-analog probing techniques, providing a more realistic understanding of metabolic activity under distinct environmental conditions.

**Importance:** Microbial communities are complex and dynamic, with different groups of microbes active under distinct conditions. Bio-Orthogonal Non-Canonical Amino Acid Tagging (BONCAT) uses synthetic amino acids to tag newly made proteins, allowing researchers to see and identify the active subset of a community. While BONCAT studies to date have used a single synthetic amino acid to evaluate cell activity in a single experimental context, here we introduce a new approach, “dual-BONCAT,” using two synthetic amino acids to track differential responses to changing conditions. After validating the approach with *E. coli*, we deployed it in a salt marsh sediment community, finding that organisms potentially feeding on plant root sugars were more active during the day, and microbes likely metabolizing sulfur were more active at night. We believe dual-BONCAT will prove useful in many studies, as it illuminates microbial community responses to changing conditions, which has important implications for ecosystem dynamics.

## Introduction

Understanding the metabolic activity of microbes in both controlled cultures and environmental communities is a major goal for microbiologists. Substrate analog probing techniques, which use analog molecular constituents to track assimilatory processes (1–3), have been adapted for use in environmental communities (4, 5). One such approach is Bio-Orthogonal Non-Canoncial Amino Acid Tagging (BONCAT), which relies on the cellular uptake of the non-canonical amino acids (ncAAs) *L*-azidohomoalanine (AHA) and *L*-homopropargylglycine (HPG), which can be incorporated into newly synthesized proteins in place of *L*-methionine without substantially affecting cell function (5, 6). AHA and HPG possess chemically modifiable azide and alkyne groups, respectively, to which fluorescently tagged molecules can be bound through “click chemistry” reactions (7, 8). Recent validation of β-ethynylserine as a threonine analog amenable to click reactions in a range of organisms offers a third ncAA for studies of anabolic activity (9).

BONCAT is therefore able to distinguish anabolically active (or more precisely, protein-synthesizing) cells from dormant cells. Downstream analysis through microscopy or sequencing (often following Fluorescence *In Situ* Hybridization, FISH, or isolation via Fluorescence-Activated Cell Sorting, FACS), enables an array of ecophysiological interpretations. For instance, BONCAT has been used to show carbon use preferences of distinct taxa in hot springs (10) and salt marsh sediment (11), active microbes in the soil microbiome (12) and biocrusts (13), and symbiotic relationships between anaerobic methane-oxidizing archaea (ANME) and sulfate-reducing bacteria in the deep sea (5).

Despite the substantial advances made possible by BONCAT in the study of cultured organisms and environmental microbial communities, all known applications of BONCAT to date have used only one ncAA in a given experiment. While this technique has allowed researchers to access the anabolically active subpopulation of a microbial community in a given experimental context, it is not capable of identifying activity patterns among a single population under different or changing conditions. Here, we present a novel approach making use of two ncAAs – AHA and HPG – in a single experiment, a technique we refer to as “dual-BONCAT”. This method offers additional versatility for more complex and environmentally representative experimental designs: by selectively labeling two distinct protein subsets, dual-BONCAT provides multiplexing capabilities that enable the labeling and analysis of two protein populations within the same cell or sample. Critically, such experiments would not require parallel incubations that may be subject to small but significant differences in community composition and physicochemical parameters, or may be unobtainable with *in situ* or sample-limited studies. With dual-BONCAT, critical spatial relationships that may dictate differential activity under distinct conditions can be retained and explored. In this study, we first validate the dual-BONCAT approach with *E. coli* cell cultures and an environmental microbial community, and then present a proof-of-concept study differentiating salt marsh sediment communities that are active during the day or night.

## Results & Discussion

### Reagent & Sample Selection

For our two ncAA-dye combinations, we selected the AHA-DBCO dye strain-promoted click reaction and the HPG-Azide dye Cu(I)-catalyzed cycloaddition reaction. These selections minimized the potential for false positive signal that could result from using the HPG-Azide dye and AHA-Alkyne dye pairings: the Cu-catalyzed “click” reaction links azide and alkyne groups, meaning unbound azide and alkyne dyes could react with each other and may be more difficult to wash away. The DBCO dye molecule does not contain a reactive alkyne group and would thus not be susceptible to such dye-dye cross-reactions. (The potential for false negative signal is considered below.)

We performed our optimization and experimental procedures on cell cultures (*Escherichia coli*) as well as an environmental system hosting a complex microbial community (surface sediment from the Little Sippewissett Salt Marsh on Cape Cod, MA). In this way, we show the efficacy of the dual-BONCAT technique in a well-constrained pure culture context, while also demonstrating the relevance of the technique (as well as the associated challenges) when used with environmental microbiomes.

### Dual-BONCAT Validation with *E. coli* Cultures

To evaluate the effectiveness of various ncAA and dye concentrations for subsequent use in dual-BONCAT experiments, we first conducted a range of mono-BONCAT tests using a single ncAA-dye pairing. Based on previously published research in both cultures and environmental systems (5, 14, 15), we tested a range of ncAA (0-50 µM HPG; 0-500 µM AHA) and dye (0.1-100 µM for both Azide-Cy3 and DBCO-Cy5) concentrations. In *E. coli* cultures, we found that the fluorescence signals from ncAA concentrations ≥5 µM and dye concentrations ≥1 µM were distinguishable from both ncAA-free and cross-reaction (i.e., AHA-Azide-Cy3 and HPG-DBCO-Cy5) control conditions under our chosen microscope settings (Figs. S1-S3). ncAA concentrations of at least 5 µM and dye concentrations of 25 or 100 µM produced particularly strong signals with median fluorescence intensities 12-64 times greater than the corresponding ncAA-free controls (Fig. S3).

To validate the dual-BONCAT approach, we conducted a series of experiments with *E. coli* cultures using 5 µM HPG, 5 µM AHA, 25 µM Azide-Cy3, and 25 µM DBCO-Cy5, based on our mono-BONCAT results. (Some conditions of higher ncAA and dye concentrations produced marginally higher fluorescence signals, but we prioritized the lower concentrations to minimize costs and avoid potential toxic effects of higher ncAA concentrations (16, 17).) 100 µL of stationary phase culture was added to 9.9 mL of sterile M9 medium; the first ncAA (ncAA#1) was added at inoculation (0h) and the second (ncAA#2), when relevant, was added after 8h of growth at 37 °C. Cultures were fixed after 24h of incubation. We tested several different experimental configurations to determine whether the fluorescence signals from the two different ncAAs were equivalent, and whether the order of ncAA addition and/or the order of post-fixation dye reactions affected the fluorescence intensities.

Overall, fluorescence signal from both ncAAs was clearly distinguishable from background and non-specific binding effects regardless of the timing and order of ncAA addition and the order of the dye reactions (Figs. S4-S5, 1). However, we broadly observed higher fluorescence when the HPG-Azide-Cy3 reaction was performed first, as the median Cy3 fluorescence was 32% brighter (p < 0.001) and the median Cy5 fluorescence was 151% brighter (p < 0.001) relative to when the AHA-DBCO-Cy5 was performed first (Fig. 1). This trend included higher intensities associated with non-specific binding effects (e.g., cases where the corresponding ncAA was absent, as in ncAA-free and mono-BONCAT cross-reaction controls), but the overall ability to distinguish true positives from false positives was largely unaffected by the order of the dye reactions: false positives could only explain 1.6% of the true positive (e.g. mono-labelled) data, at most, regardless of dye order. Initially, we had hypothesized that false negatives could result from performing the HPG-Azide reaction first, as the alkyne-azide click reaction could in theory link AHA azide groups with HPG alkyne groups, thereby blocking dye molecules from binding to their corresponding ncAAs and diminishing overall signals. The fact that we did not observe a decrease in fluorescent signal when the HPG-Azide-Cy3 reaction was done first suggests that the AHA-HPG reaction is not favorable – perhaps due to steric hindrance associated with peptide chains – and is not a substantial concern when using the dual-BONCAT approach. Therefore, we recommend performing the HPG-Azide reaction first, and report the results with this dye order unless otherwise specified. Compared with the corresponding mono-BONCAT (mono-stained) controls, the median fluorescence signal of HPG-labeled cells increased (45% increase, p < 0.001) when both dye reactions were performed. In contrast, the AHA-associated signal decreased by 48% (p < 0.001).

**Fig. 1:**
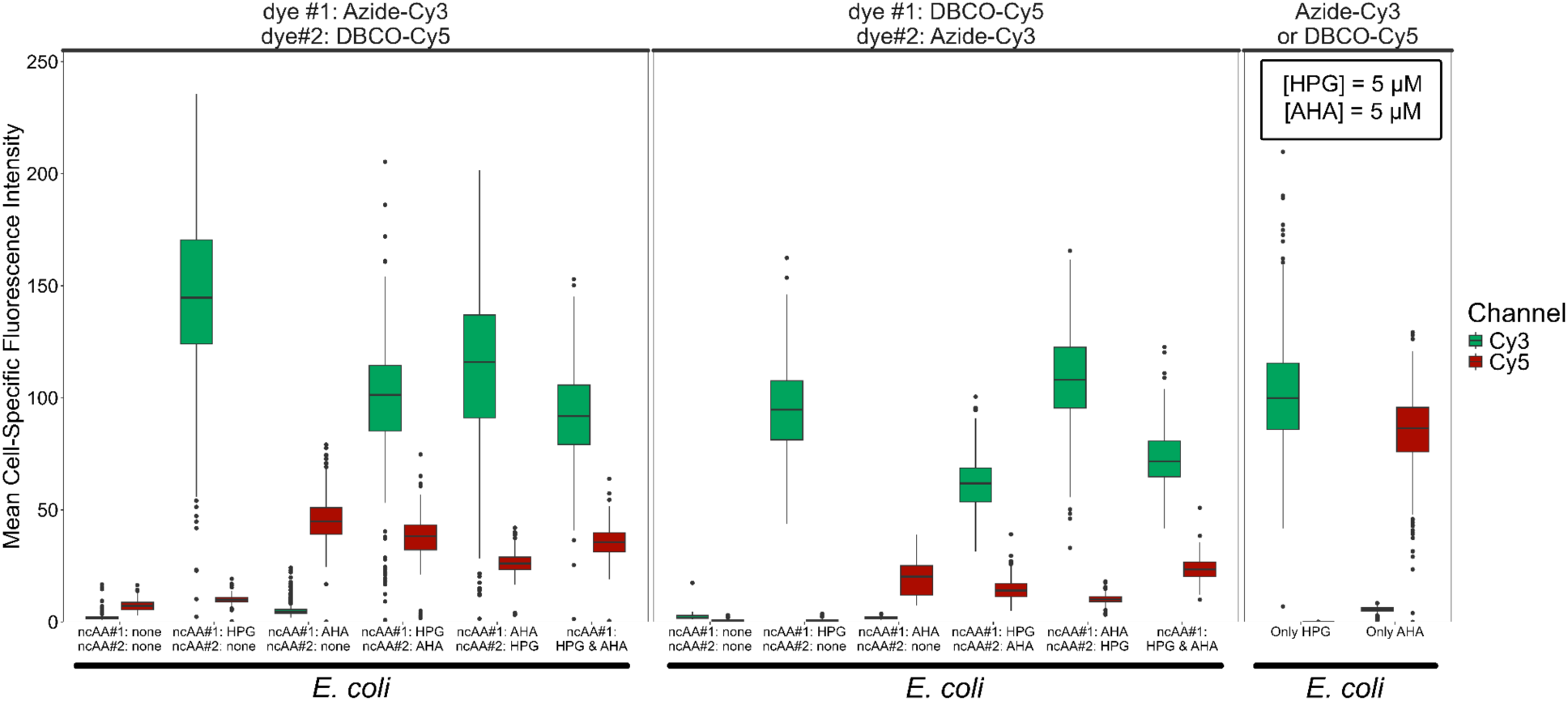
Boxplots of mean fluorescence intensity values of cells in dual-BONCAT experiments with *E. coli* cultures. Tests varying the sequence of ncAA addition (see column labels along x-axis) were performed, with the Azide-Cy3 click reaction conducted first (left panel) and with the DBCO-Cy5 click reaction conducted first (middle panel). The procedural control in which *E. coli* cells were mono-labeled and mono-stained is shown in the right panel.

In comparing the order of the ncAA addition, Cy3 fluorescence was 13% dimmer (p < 0.001) but the Cy5 fluorescence was 47% brighter (p < 0.001) when HPG was added first relative to when AHA was added first (when the HPG-Azide-Cy3 dye reaction was done first). This trend was repeated when the dye order was switched. This observation that a given ncAA generates brighter signal when added second rather than first is intriguing, and may be due to protein turnover dynamics whereby proteins synthesized earlier break down over time. Compared to the mono-BONCAT (dual-stained) controls, the median fluorescence signal of Cy3 and Cy5 decreased by 36% (p < 0.001) and 21% (p < 0.001), respectively, when both ncAA were added simultaneously.

The median HPG-Azide-Cy3 signal in the mono-labeled, dual-dyed experiment was significantly brighter (by 222%, p < 0.001) than the AHA-DBCO-Cy5 (mono-labeled, dual-dyed) signal. This relationship was also observed when both ncAAs were added at different times: HPG-Azide-Cy3 signal was 210% (p < 0.001) brighter when the Azide-Cy3 dye reaction was performed first, and 462% (p < 0.001) brighter when it was performed second (when both orders of ncAA addition were taken into account, in both cases) (Fig. 1). Since fluorescence intensity profiles of mono-BONCAT experiments were broadly similar between HPG and AHA (Fig. S3), these findings suggest that the dynamics change when both ncAAs are present: in such cases, HPG may be more effective than AHA at integrating into newly synthesized proteins, and/or the HPG-Azide click reaction may be favored over the AHA-DBCO reaction.

To determine if dual-BONCAT is capable of distinguishing distinct populations that were anabolically active at different stages of an incubation, we looked at “differential labeling” patterns, where some cells were only labeled with Azide-Cy3, some were only labeled with DBCO-Cy5, and some were labeled with both dyes. Because the intensity of BONCAT labelling cannot always be taken as a quantitative indication of anabolic activity (due to, among other factors, distinct proteome compositions, varying rates of ncAA:Met substitutions, and variability in the rate of dye entry into the cell across different species (17–19)), we maintained a binary, “labeled” or “not labeled” approach to interrogate differential labeling. Threshold fluorescence intensity values were set by the maximum intensity, in either channel, of the condition in which no ncAAs were added but dye reactions were performed. Cells whose mean intensity exceeded these values in a given fluorescence channel were considered “labeled” with the corresponding ncAA.

We first considered the condition in which both ncAAs were present at the same concentration and for the same duration, reasoning that, in this case, no differential labeling should occur. Indeed, all analyzed cells were dual-labled (Fig. S6, S7). Next, we investigated differential labeling patterns among *E. coli* cells when the ncAAs were added at different times. Single cell-level observations showed clear visual evidence of differential labeling: some cells were exclusively labeled with Azide-Cy3, some were exclusively labeled with DBCO-Cy5, and some were labeled with both fluorescent molecules (Fig. 2). Some cells were labeled with neither dye, suggesting they were anabolically inactive during the 24-hour culturing experiment, potentially representing quiescent or dead biomass carried over from the inoculum. Using the threshold-based assessment of labeling, we confirmed that differentially labeled subpopulations were present (Fig. S7). These observations confirm the ability of dual-BONCAT to detect distinct subpopulations that are anabolically active during different phases of an experiment. The fact that the majority of cells in these 24-hour experiments were dual-labeled even when the second ncAA was added eight hours after the first is not surprising given the dynamics of exponential growth in pure culture. Different timelines of adding the ncAAs and terminating the experiment could likely be calibrated to attain a larger proportion of differentially labeled cells.

**Fig. 2:**
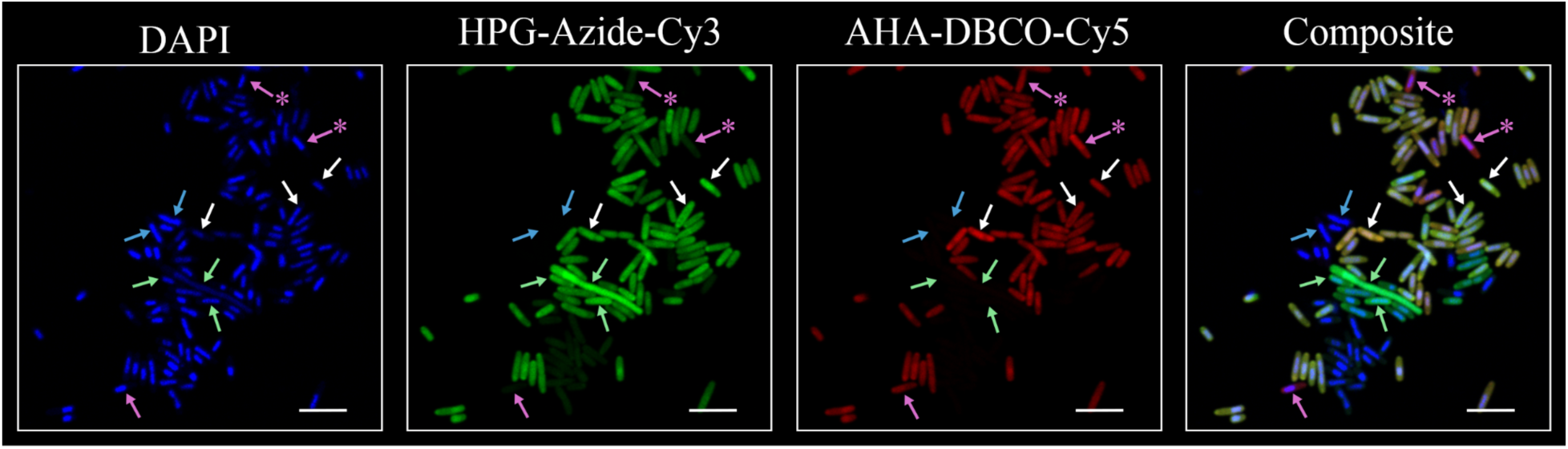
Fluorescence microscopy field of view of a dual-BONCAT experiment with *E. coli* cells. HPG was added at the start of the experiment; AHA was added 8 hours later, and the experiment was stopped after 24 hours. The Azide-Cy3 click reaction was performed first. Arrows indicate select cells representative of the distinct subpopulations identifiable with dual-BONCAT. Blue arrows: cells that were not labeled with either ncAA. White arrows: cells that incorporated both ncAAs. Green arrows: cells that only incorporated HPG. Magenta arrows: cells that only incorporated AHA. Magenta arrows with asterisks indicate cells with very low Cy3:Cy5 ratios (see Fig. S7) that dominantly incorporated AHA, but whose Cy3 intensities were slightly above the threshold value set for our binary “labelled” designation. Scale bar indicates 5 µm.

### Dual-BONCAT in an Environmental Microbial Community

Dual-BONCAT was next deployed with an environmental microbial community, using surface sediment from Little Sippwissett Salt Marsh, which contains a dense and taxonomically and metabolically diverse microbiome (11, 20–23). Our aim was to observe and ultimately characterize the subpopulations of cells that were 1) anabolically active during the 12-hour sunlit day, 2) anabolically active during the 12-hour dark night, 3) anabolically active during both the day and night periods, and 4) not active during the 24-hour experiment. Sediment was recovered from the marsh, returned to the lab, and distributed to mesocosm incubations within several hours. In all experimental conditions, one ncAA was added at dawn (ncAA#1) and the incubation was placed in sunlit conditions; 12 hours later, the incubation was moved to a dark incubator and the second ncAA (ncAA#2) was added. The temperature was maintained at approximately 25 ℃ throughout.

Three ncAA concentration profiles were tested: 5 µM HPG & 5 µM AHA & (referred to as 5/5 below); 50 µM HPG & 50 µM AHA (50/50); and 50 µM HPG & 500 µM AHA (50/500). These conditions expand beyond the ncAA concentrations optimized for *E. coli* (5 µM for each ncAA) in order to match HPG concentrations previously optimized for LSSM sediment (11, 22), and to account for the fact that AHA is sensitive to reduction under alkaline sulfidic conditions such as those in Little Sippewissett salt marsh sediments (14), meaning that its “felt” concentration was likely lower than what was added. All dye concentrations were maintained at 25 µM based on the *E. coli* optimization experiments described above.

While fluorescence signal derived from HPG incorporation was substantially brighter than that from AHA incorporation across all conditions when their concentrations were equal, both ncAAs were statistically distinguishable from background and non-specific binding effects under elevated ncAA conditions (e.g., the 50/50 and 50/500 experiments; Figs. 3; S8-S9). (The p-values were < 0.001 for all ncAA combinations except for the Cy5 signal when both ncAAs were added at the start of the 50/50 experiment, p = 0.052, though cells brighter than the non-specific binding signal were detected.) The 5/5 condition resulted in fluorescence signals that were largely indistinguishable from background and non-specific binding signals (Fig. S10). Prior to click-chemistry reactions, cells from the LSSM sediment incubations were extracted to separate them from the mineral matrix. Compared with *E. coli* culture experiments run under analogous ncAA concentrations, median fluorescence intensity of *E. coli* cells subjected to the cell extraction process decreased by 18% (p < 0.001) and 54% (p < 0.001) for Cy3 and Cy5, respectively, indicating that the extraction approach itself compromised signal intensity (but not detectability compared with control treatments). Under the 50/50 condition, the median fluorescence signal of labeled sediment cells decreased by 47% (p < 0.001) for Cy3 and 54% (p < 0.001) for Cy5 when both ncAAs were added simultaneously, compared with their mono-BONCAT (dual-stained) controls (Fig. 3A). In contrast, Cy3 increased by 27% (not statistically significant, p = 0.34) and Cy5 increased by 106% (p < 0.001) in the 50/500 experiment (Fig. 3B).

**Fig. 3:**
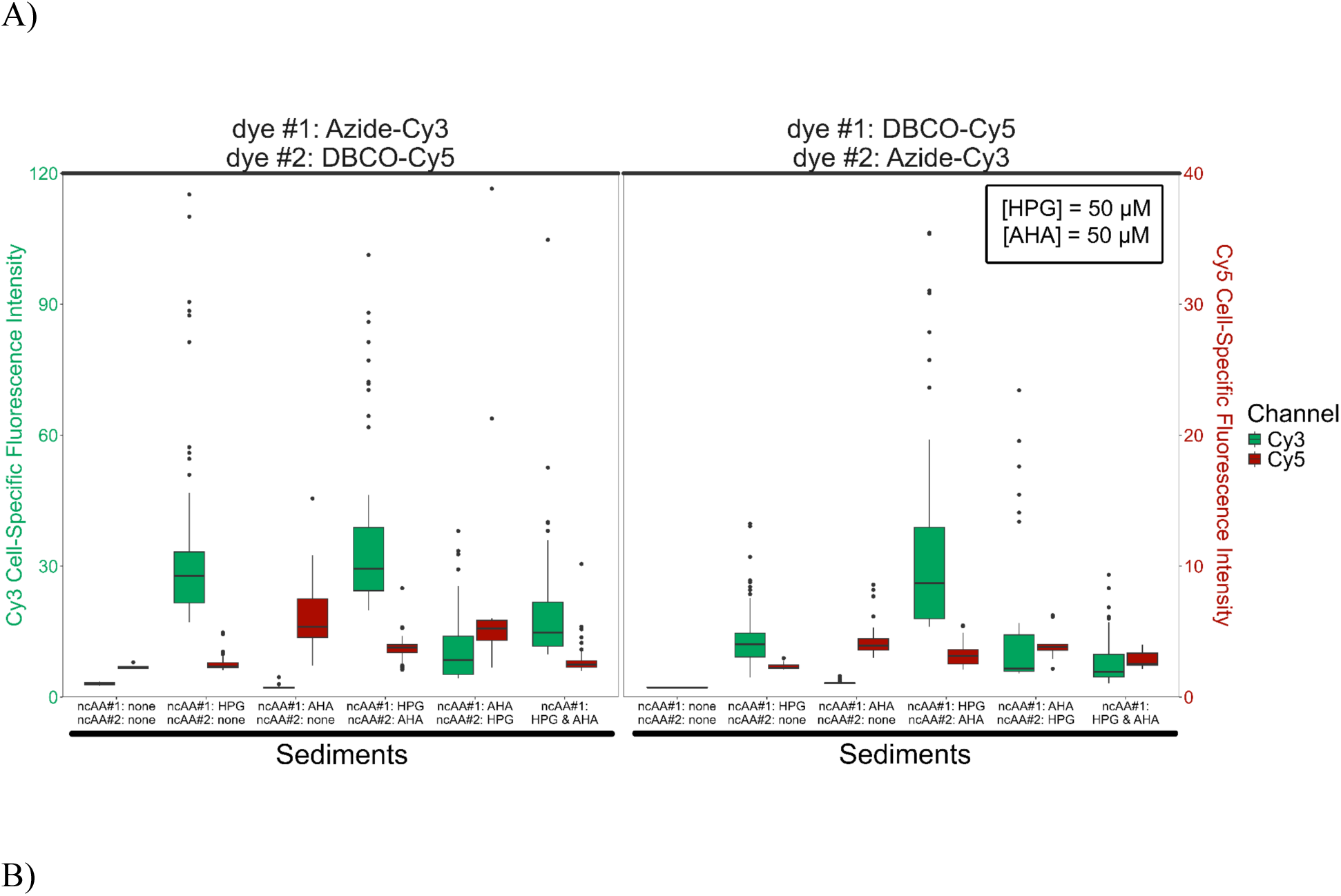
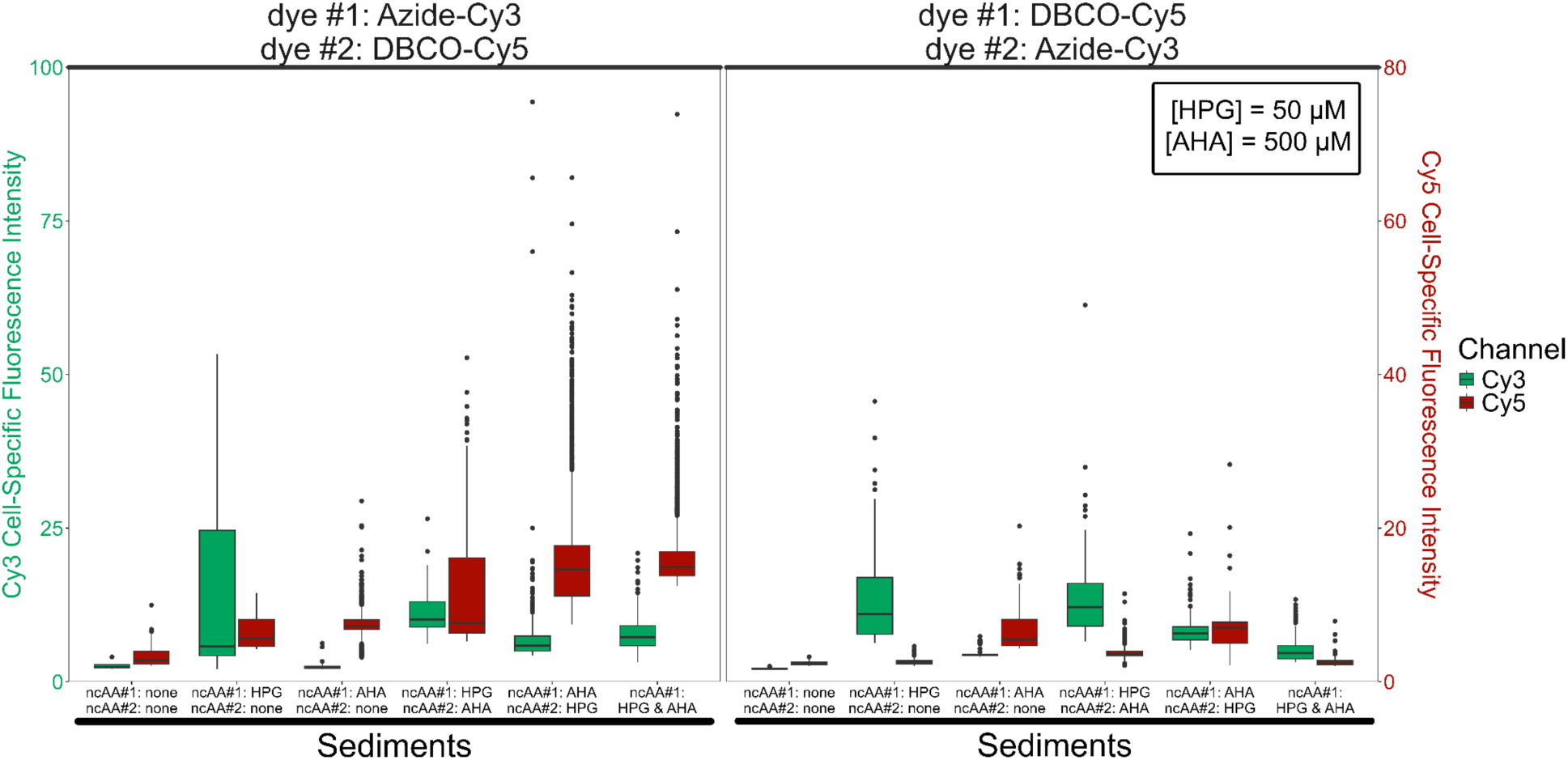
Boxplots of mean fluorescence intensity values of cells in dual-BONCAT experiments with extracted salt marsh sediment cells. Tests varying the sequence of ncAA addition (see column labels along x-axis) were performed, with the Azide-Cy3 click reaction conducted first (left panel) and with the DBCO-Cy5 click reaction conducted first (middle panel). A) Data from the 50 µM HPG & 50 µM AHA condition; B) data from the 50 µM HPG & 500 µM AHA condition.

In comparing the order of ncAA addition for the 50/50 experiment, Cy3 fluorescence was 248% brighter (p < 0.001) but the Cy5 fluorescence was 28% dimmer (p < 0.05) when HPG was added first relative to when AHA was added first (and HPG-Azide-Cy3 dye reaction was performed first in both cases). This trend was repeated when the dye order was switched, and, in contrast to the *E. coli* experiments, indicates that a given ncAA generates brighter signal when added first, potentially due to slower protein turnover or degradation in the environmental community compared with a pure culture.

In all four dual-BONCAT permutations (testing the order of ncAA additions and dye reactions) with the 50/50 experiment, the HPG-Azide-Cy3 signal was generally brighter than the AHA-DBCO-Cy5 signal. This was also observed when both ncAAs were added: HPG-Azide-Cy3 signal was 432% (p < 0.001) brighter when the Azide-Cy3 dye reaction was performed first, and 166% (p < 0.001) brighter when it was performed second. These patterns are broadly improved (i.e., the AHA-DBCO-Cy5 signal is generally brighter) under the 50/500 condition, indicating that relative concentrations of the ncAAs, even under sulfidic conditions, can be used to “tune” detectability and relative signal intensities.

Experiments in which AHA was added first (at sunrise) and HPG was added second (at night) resulted in a greater overall separation of fluorescent signal compared with control conditions, offering the best opportunity to detect cells growing at different times.

We next considered differential labeling patterns at the single-cell level. When both ncAAs were added at the start of the experiment, nearly all cells were dual-labeled (Fig. S11). When one ncAA was added at sunrise and the other at sunset, we observed distinct subpopulations of cells exhibiting only HPG-Azide-Cy3, only AHA-DBCO-Cy5, both dyes, and neither dye (Fig. 4).

**Fig. 4:**
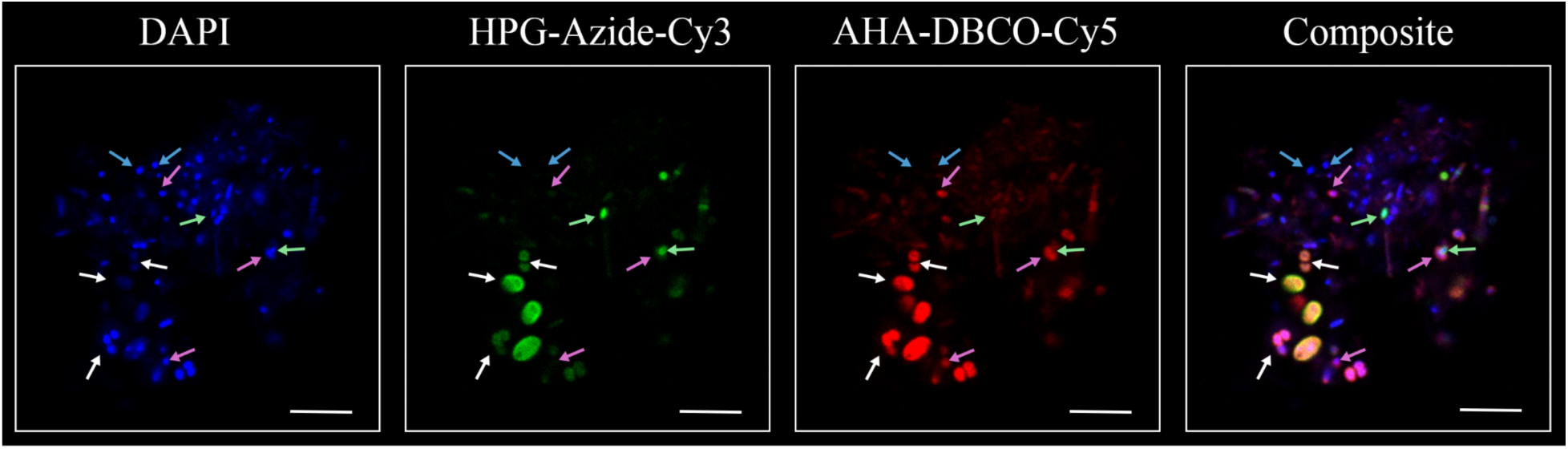
Fluorescence microscopy field of view of a dual-BONCAT experiment with extracted salt marsh sediment cells. 50 µM HPG was added at the start of the experiment; 500 µM AHA was added 12 hours later, and the experiment was stopped after a single 24-hour day-night cycle. The Azide-Cy3 click reaction was performed first. Arrows indicate select cells representative of the distinct subpopulations identifiable with dual-BONCAT. Blue arrows: cells that were not labeled with either ncAA. White arrows: cells that incorporated both ncAAs. Green arrows: cells that only incorporated HPG. Magenta arrows: cells that only incorporated AHA. Scale bar indicates 5 µm.

### Interpretation of Dual-BONCAT Subpopulations

Interpreting the physiological / ecological meaning of the three differentially labeled cell populations is not entirely unambiguous. Dual-BONCAT experiments can be generalized as follows: one ncAA (ncAA#1) is added at an initial time point (T_1_), a second ncAA (ncAA#2) is added at a subsequent time point (T_2_), and the experiment is ended (i.e., all metabolic activity is ceased) at a later, final time point (T_F_); see Fig. S12. The three differentially BONCAT-labeled populations can be identified as the cells exhibiting fluorescence associated with only the first ncAA (ncAA#1), both ncAAs (ncAA#1&2), and only the second ncAA (ncAA#2). We propose the following interpretations of these three populations:

– The ncAA#1 population is only active between T_1_ and T_2_.
– The ncAA#1&2 population includes cells that either are active from both T_1_ to T_2_ *and* T_2_ to T_F_, or are active between T_2_ and T_F_. Because ncAA#1 could be present throughout the experiment, it could still be used by organisms exclusively active after T_2_. Such dual-labeled cells may exhibit a particularly high affinity for ncAA#1.
– The ncAA#2 population is only active between T_2_ and T_F_. We suggest that the lack of ncAA#1 incorporation in this population could be due to the relative ncAA concentrations during the second phase of the experiment, after the ncAA#1 pool has been depleted. If these cells were anabolically active when only ncAA#1 was present, it would have been incorporated.

These interpretations are predicated on previous observations that all taxa queried to date are capable of using each of the two ncAAs at roughly equivalent incorporation frequencies ((17) and references therein). If these frequencies differ between the two ncAAs, further calibration of concentrations would be needed to attain similar fluorescence signals. (For reference, HPG and AHA replace an estimated 1 out of every 500 and 390 methionine residues in newly synthesized proteins, respectively (24), while the newly reported THRONCAT approach has a substitution efficiency of 1:40 (9).)

### Dual-BONCAT-FACS Reveals Day– and Night-Active Taxa in Salt Marsh Sediment

To demonstrate the utility of the Dual-BONCAT technique for studying the dynamics of environmental microbial communities, we sought to identify which microbes in salt marsh sediment were active (synthesizing proteins) during sunlit daytime hours and which were active during nighttime darkness. Critically, this approach did not require separate incubations, which obviates concerns about distinct microbial communities or physicochemical conditions across supposed “replicate” samples.

Two lab-based experimental incubations of shallow (0-3 cm) salt marsh sediment were performed. In Condition 1, HPG was added at sunrise (T_1_) and AHA was added 12 hours later (T_2_), when the incubation was moved to a dark environment. In Condition 2, the order of ncAA addition was reversed. We also included a negative control treatment that did not receive any ncAAs but did receive both dyes. The downstream use of Fluorescence-Activated Cell Sorting (FACS) requires stained cells to be in suspension rather than affixed to slides, prompting some modifications to the slide-based protocol (detailed in the Materials & Methods section). In particular, we found that 25 µM Azide-Cy3 and 1 µM DBCO-Cy5, when applied to 50 µM concentrations of both ncAAs, was sufficient to differentiate true positive from false positive signal in cell sorting experiments. For both experimental conditions, we confirmed the detection of all three populations of cells (ncAA#1, ncAA#1&2, and ncAA#2) under the microscope prior to cell sorting.

The three BONCAT-active populations from each of the two incubations were isolated using FACS, based on gates determined using negative and mono-BONCAT positive controls (Fig. S13). With the exception of the ncAA#2 population (active only at night) from Condition 1, which generated FACS cell counts below the detection limit, between 2.1×10^4^ and 7.2×10^5^ cells were sorted for each population (Table S1). Subsequent high-throughput 16S rRNA gene amplicon sequencing allowed us to compare these distinct populations and infer ecophysiological differences among day– and night-active taxa.

Among the taxa that were most prominent within the day-active (ncAA#1) population (Fig. 5; Tables S2, S3), the *Beijerinckiaceae* genus *Methylobacterium* is commonly associated with plants, and can use one-carbon compounds, including methanol in plant exudate, as a carbon source (25, 26). At LSSM, roots of the cordgrass *Spartina alterniflora* permeate the sediment and release a chemically diverse array of organic molecules. Heterotrophs feeding on exudate, such as *Methylobacterium*, were likely stimulated by the *Spartina*’s daytime photosynthetic activity; a similar relationship was observed with nitrogen-fixing microorganisms surrounding *Spartina* roots, whose activity was substantially enhanced within an hour of light exposure (27).

**Fig. 5:**
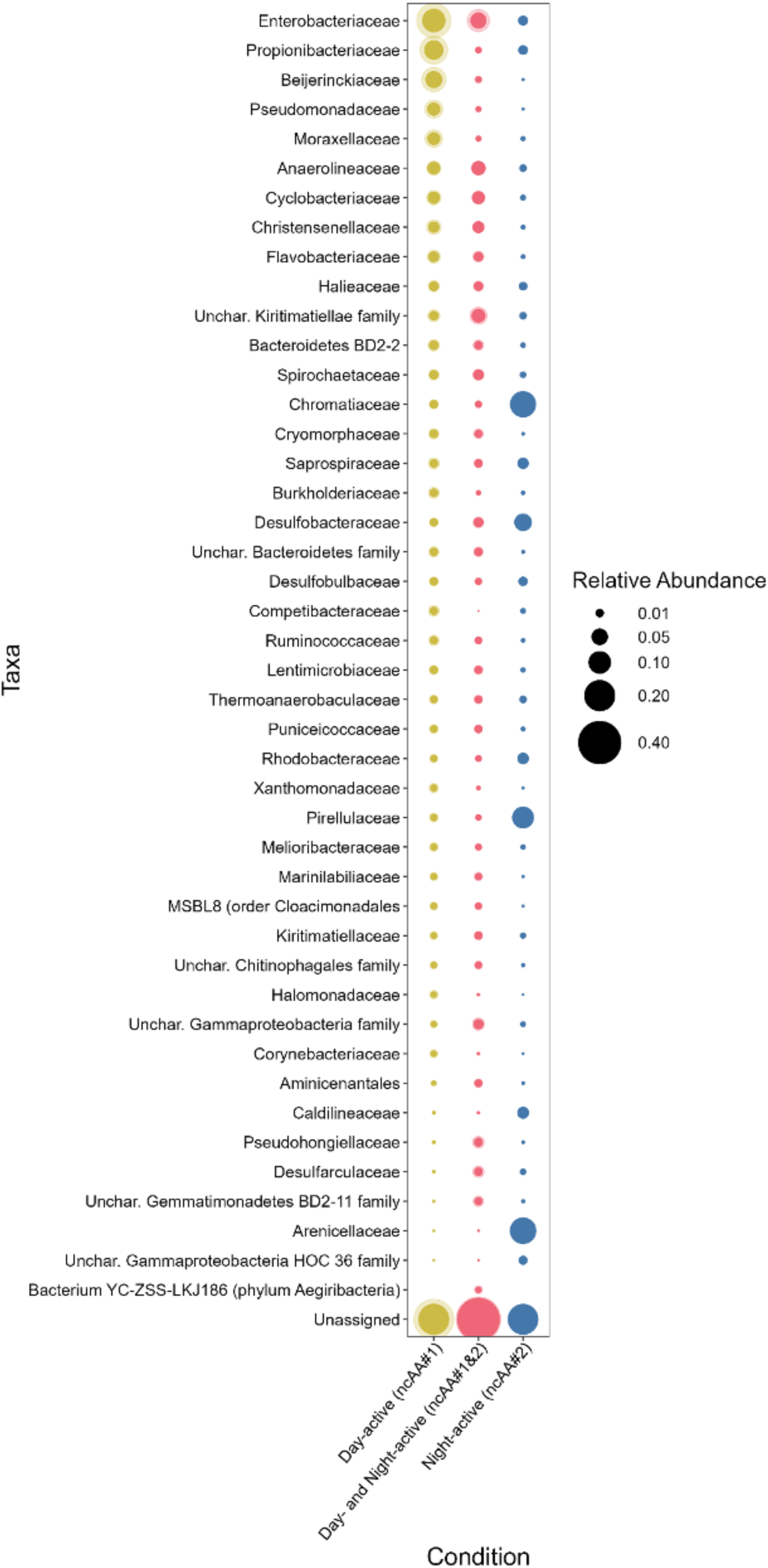
Relative abundances of family-level taxonomic groupings derived from dual-BONCAT-FACS-16S analyses of LSSM sediments. Each row corresponds to a family that comprised at least 1% of the overall relative abundance in at least one of the five sorted and sequenced fractions. Each column and color corresponds to a given ncAA condition: yellow indicates cells that incorporated only ncAA#1 (regardless of which ncAA was added first), red indicates cells that incorporated both ncAAs (regardless of which was added first), and blue indicates cells that incorporated only ncAA#2 (when HPG was added second; the other condition did not result in sufficient cells for sequencing). The size of each circle indicates the mean relative abundance for a given family and ncAA condition; the faint “halo” around each data point conveys the standard deviation when two communities were sequenced for a given condition. See Table S2 for the data used to generate this figure.

Sulfur-cycling taxa were highly abundant members of the night-active community (ncAA#2 population). *Chromatiaceae*, the purple sulfur bacteria that comprise a key constituent of the syntrophic “pink berry” consortia (21), accounted for 14% of the night-active population; while this taxon is photosynthetic, its dark-phase respiration metabolism (28, 29) may be more tightly coupled with protein synthesis. (We are confident that this enrichment was not due to *Chromatiaceae* autofluorescence because signals in ncAA-free control experiments did not rise above the threshold of “positive” detection; Fig. 3). *Arenicellaceae*, whose close relative UBA868 may play a foundational role in marine sulfur oxidation (30), and the sulfate-reducing bacteria family *Desulfobacteraceae* were both preferentially active at night, perhaps responding more favorably than other anaerobic taxa to expanding anoxic zones that accompany nighttime decreases in oxygenic photosynthesis (31). Other abundant taxa, such as the heterotrophic *Anaerolineaceae* and *Cyclobacteriaceae*, did not exhibit night-specific activity patterns; these organisms likely operate independent of diurnal cycles and depend more on ever-present organic matter rather than shifting redox stratification or specific molecules provisioned during light or dark metabolic phases.

## Conclusion

Dual-BONCAT offers a new capability for the visualization and identification of differentially active subpopulations within cell cultures and environmental microbiomes. By revealing organisms that are anabolically active under distinct conditions within the same physical environment, novel syntrophic or antagonistic relationships can be queried, and a more realistic understanding of the condition-specific activity of microbial constituents can be attained. We found that, in *E. coli* cultures and a salt marsh sediment microbial community, both AHA and HPG uptake was detectable, and that distinct subpopulations of cells active during different phases of an experiment were distinguishable. The order of ncAA addition did not have a consistent effect on fluorescent intensity, but the order of dye reaction did: overall, performing the HPG-azide click reaction prior to the AHA-DBCO reaction yielded brighter signals. Downstream FACS and high-throughput 16S rRNA gene sequencing of distinct labeled subpopulations allowed for the ecophysiological interpretation of taxa active during the day and during the night, all without disrupting the community or relying on pseudoreplicate experiments.

Challenges associated with dual-BONCAT are similar to those associated with mono-BONCAT, and largely relate to optimizing signal-to-noise ratios in environmental microbiomes. Low signal challenges may be accentuated when both ncAAs are present, since competition for Met substitution would be expected to lower incorporation of each ncAA by approximately half. We recommend that users customize ncAA concentrations, dye concentrations, and imaging parameters to optimize for their specific applications, as diffusivity and non-specific binding effects can vary with biofilm density, mineralogical context, and other physicochemical factors. However, given the broad flexibility of mono-BONCAT more generally, we anticipate that the methods and parameters we describe here will be a successful starting point for most applications, including isolate cultures and complex environmental samples.

We envision many promising use cases for the dual-BONCAT approach, which allows researchers to interrogate how a specific microbial community responds to changing conditions. For example, the technique could track spatial patterns of protein synthesis within cell aggregates or biofilms, or identify anabolically active taxa, in response to a shift in temperature, pH, redox potential, predation, oxygen concentration, or any other manipulatable variable. Protein pull-down experiments, which capture newly-synthesized proteins for downstream sequencing (32), could incorporate dual-BONCAT to identify sub-proteomes linked with responses to environmental challenges. Because of microscale heterogeneity in microbial community composition (33, 34), “replicate” experiments are often insufficient in these contexts. Using dual-BONCAT to observe the response of a single community as conditions change offers a more realistic, representative view of dynamic microbiomes.

## Materials and Methods

### Sample Growth & Collection

*E. coli* was used first to identify the optimal concentrations of AHA and HPG, along with their conjugate dye concentrations, and second to test dual-BONCAT approaches. The selection of *E. coli* as our experimental organism was based on its rapid replication time, high expression capabilities, and previous utility in BONCAT method development. Given that both non-canonical amino acids used in this study, AHA and HPG (both supplied by Click Chemistry Tools, Scottsdale, AZ), serve as surrogates for methionine, M9 medium, a growth medium that lacks methionine, was used to grow the cultures. *E. coli* was initially inoculated from a glycerol stock into M9 medium and grown overnight. (This *E. coli* stock had been grown in LB nutrient-rich medium prior to being stored in glycerol.) We then took 100 µL of early stationary phase *E. coli* culture in M9 to inoculate 9.9 mL of sterile M9 media. The first ncAA was added at the time of inoculation; the second, when relevant, was added at a designated intermediate time point (8 hours). All ncAA and dye concentrations are indicated in the main text for each specific experiment.

To explore the use of Dual-BONCAT with environmental microbial communities, we used sediment from Little Sippewissett Salt Marsh on Cape Cod, MA, USA. Sediment from 0-3 cm depth was collected from the “Berry Pool” (located at 41.5758° latitude, –70.6394° longitude); these sediments contained small *Spartina alternaflora* grasses and are marked by sharp redox gradients with diurnal geochemical shifts, as well as a strong heritage of microbial diversity studies, some of which have successfully deployed BONCAT and BONCAT-FACS-16S techniques (11, 20, 22, 31). Immediately following collection, sediment was returned to the lab and placed next to a large window at room temperature. Incubation experiments began within two days after collection, and ncAAs were added at designated time points from stock solutions to reach the desired final concentrations. AHA is sensitive to reduction under alkaline sulfidic conditions such as those in LSSM sediments (14); thus, the AHA added to the incubations should be viewed as an upper bound on the “felt concentration” within the experiments. Incubations were conducted in 15 mL Falcon tubes with 5 mL of sediment and 5 mL of co-localized marsh water. For day/night dual-BONCAT experiments, ncAA#1 was added at sunrise, and incubations were placed next to a large window with direct daytime light. Twelve hours later, ncAA#2 was added and the incubations were placed in a sealed incubator at room temperature. Controls with no ncAAs, only one ncAA, and both ncAAs added at the start of the experiment were also included. For fluorescence activated cell sorting experiments, control incubations were set up for 1) a Cy3 Positive Control (mono-BONCAT with HPG-Azide-Cy3), 2) a Cy5 Positive Control (mono-BONCAT with AHA-DBCO-Cy5), 3) a Cy3&Cy5 Positive Control (both ncAAs added for the full duration of an incubation experiment), and 4) a Negative Control (neither ncAA was added, but Azide-Cy3 and DBCO-Cy5 dyes were). All ncAA and dye concentrations are specified in the text.

To stop the incubations at the designated time points, samples were chemically fixed to cease metabolic activity and prepare the cells for click chemistry reactions and imaging. To do this, 0.5 mL aliquots of culture or sediment slurry were placed in sterile 2 mL tubes, and 1.5 mL of 0.22-µm filter sterilized 3% paraformaldehyde in 1x PBS was added to each tube. This mixture was mixed briefly by vortex and kept at room temperature for one hour. The tube was then centrifuged at 8,000 x *g* for 5 minutes and the supernatant was discarded. The pellet was washed twice with 1x PBS (by resuspending in 1 mL of 1x PBS, centrifuging at 8,000 x *g* for 5 minutes, and removing the supernatant) and ultimately resuspended in 1 mL of 1:1 PBS:ethanol. The sample was then kept at 4 °C until downstream processing.

### Cell Extraction from Sediment Samples

To recover cells from LSSM sediment and minimize the effects of autofluorescent mineral grains, a cell extraction was performed following a protocol established by Couradeau et al. for the soil microbiome (12) and successfully used in LSSM sediment (11). Fixed sediment was centrifuged at 5,000 x *g* for 3 minutes, the supernatant was removed, and ∼250 µL of sediment (or fixed culture, for controls) was transferred to a new 2 mL tube. 1450 µL of sterile 1x PBS and 34.7 µL of sterile 1% Tween20 were added to the tube, which was then inverted several times and vortexed at maximum speed for 5 minutes to thoroughly mix the sample and promote the removal of cells from sediment grains. Tubes were then centrifuged at 500 x *g* for 55 minutes to separate cells (into the supernatant) from sediment (which formed a pellet). The supernatant was then transferred to a new 2 mL tube, being sure to avoid any sediment.

A second extraction of the initial sediment was then performed by adding 870 µL of sterile 1x PBS and 20.8 µL of sterile 1% Tween20 to the tube containing the sediment. The tube was inverted several times, vortexed at maximum speed for 5 minutes, and centrifuged at 500 x *g* for 35 minutes. The supernatant was then recovered and pooled with the previously collected supernatant from the first extraction. The pooled extraction product was then centrifuged at 16,000 x *g* for 6 minutes to pellet the cells. The supernatant was discarded and the pellet was resuspended in 250 µL of 1:1 PBS:ethanol. These fixed, extracted cells were then used for downstream cell staining analyses.

### Cell Staining with Click Chemistry for Microscopy

10 µL of fixed cells were added to a single well of a slide (Tekdon slides 8-82 with Teflon and poly-L-lysine coating) and dried at room temperature. To dehydrate and permeabilize the cells, 10 µL of 50%, 80%, and 96% ethanol were sequentially added to each well for at least 3 minutes each, and dried at room temperature in between. (A gentle stream of filtered pressurized air can be used to accelerate the drying process.) We found that performing the dehydration and click chemistry reactions directly on the wells rather than submerging the entire slide in coplin jars reduced cross-contamination between wells.

The click chemistry reactions described below can be done either in isolation (e.g., AHA-DBCO or HPG-azide for a “mono-BONCAT” application) or in series for dual-BONCAT. While we initially thought performing the AHA-DBCO reaction first would be best in order to avoid potential false negatives from AHA azides reacting with HPG alkynes, assessment of the resulting signal suggests this effect may be negligible. Instead, we recommend performing the HPG-Azide reaction first as this order of operations retained a brighter Cy5 fluorescence from the AHA-DBCO signal.

#### HPG-Azide Reaction

To perform the Cu(I)-catalyzed HPG-azide reaction, we first made two “premix” solutions in separate 2 mL tubes according to Table S4. (All volumes mentioned here can be adjusted proportionally based on the number of samples and volume of dye mix needed to fully cover the analyte cells, as well as the desired dye concentration; the volumes listed here assume a 25 µM final dye concentration and 20 µL of dye mixture applied to each microscope slide well.) Premix 1 contained the 100 mM solutions of Na-ascorbate and aminoguanidine-HCl (both in 1x PBS; solutions made fresh on the day of use), as well as the 1x PBS. Premix 2 contained the 20 mM CuSO_4_, 50 mM Tris-HydroxyPropylTriazolylmethylAmine (THPTA), and concentrated (1 mM) azide dye (Cy3 picolyl azide dye, Click Chemistry Tools, Scottsdale, AZ); these components were left to incubate in the dark at room temperature for 3 minutes. The final dye concentration can be adjusted based on the volume and concentration of the dye stock solution used; overall volume can be maintained by adjusting the amount of 1x PBS. The two premixes were then combined and inverted once to mix gently.

The incubation of cells with the dye mixture was performed in a humid chamber consisting of a 50 mL Falcon tube placed on its side, with ultrapure water-or 1x PBS-saturated kimwipes inside. 20 µL of dye mixture was applied to each slide well, and the slide was carefully inserted into the humid chamber, making sure the lens of liquid remained intact on top of each well. The humid chamber was then capped and placed in the dark at 46 °C for 30 minutes. Following the incubation, the dye was removed by carefully pipetting the solution off from the edge of the well (so as to not disturb the cells in the center) and rinsing each well three times with 20 µL of 1x PBS (applied and removed by pipette). Finally, the slide was dehydrated with 10 uL of 50% ethanol in 1x PBS for 3 minutes and then dried under a gentle stream of pressurized air.

#### AHA-DBCO Reaction

To perform the strain-promoted AHA-DBCO reaction, 20 µL of fresh 100 mM 2-chloroacetamide (in 1x PBS) were added to each slide well and incubated in the humid chamber for 1 hour at 46°C. A dye mix was made with concentrated DBCO-Cy5 dye (Click Chemistry Tools, Scottsdale, AZ) and the chloroacetamide solution; the volume and concentration should be adjusted to attain the desired final concentration (previously recommended to be between 100 nM – 1 µM (14)). We found a final dye concentration of 25 µM to work best for *E. coli*, so we used this concentration for the salt marsh sediment samples as well. The chloroacetamide was removed and 20 µL of the DBCO-Cy5 dye mix was added to each well. Slides were left to incubate in the humid chamber for 30 minutes at 46 °C in the dark. Unbound dye was removed by adding 20 µL of 1:1 dimethyl sulfoxide:PBS to each slide well and incubating in a humid chamber at room temperature for 20 minutes. Wells were then rinsed three times with 20 uL of 1x PBS. Finally, the slide wells were dehydrated with 10 uL of 50% ethanol in 1x PBS for 3 minutes and then dried under a gentle stream of pressurized air.

#### DAPI Counterstaining

To visualize all cells – and thereby confirm that BONCAT signals corresponded with biomass while also allowing for cell abundance calculations – the general DNA stain 4’,6-diamidino-2-phenylindole (DAPI) was used. For each well, 6 µL of a 1:1 mixture of 5 ng/µL DAPI:VectaShield Antifade Mounting Medium was added and spread across each well by adding a glass coverslip.

### Fluorescence Microscopy

Microscopic analysis was performed on a Leica Stellaris 5 confocal laser scanning fluorescence microscope equipped with a white light laser and tunable excitation wavelengths and emission windows. For all imaging, a 63x immersion objective lens was used and carried out in the Leica Application Suite X (LAS-X). Imaging parameters for all dyes and both cell culture and environmental samples are provided in Table S5. All imaging parameters were kept constant for experimental and control treatments in order to provide directly comparable data sets (with the exception of the DAPI detector gain, as the sediments were overexposed when the gain was set to 8%). We also found that there was substantial reflection when the detector emission window was <15 nm away from the laser excitation, which is why we deviated from the default settings in LAS-X. For each field of view, three images – corresponding to all cells (DAPI), all AHA-incorporating cells (Cy5), and all HPG-incorporating cells (Cy3) – were acquired and analyzed in parallel.

### Image Analysis

Images were analyzed using FIJI / ImageJ version 2.14.0/1.54f (35). Composite and individual channel images were generated and brightness was adjusted uniformly using our FIJI-Multi-Channel-RGB-Stacker plugin (https://github.com/mankeldy/FIJI-Multi-Channel-RGB-Stacker) with the settings in Table S6. Cell-specific fluorescence intensity data was extracted using our FIJI-Multi-Channel-Cell-Counter plugin (https://github.com/mankeldy/FIJI-Multi-Channel-Cell-Counter) with the settings in Table S7. Initially, we noticed our DAPI images were slightly offset (roughly 0.25 µm) from our Cy3 and Cy5 images, so we used the MultiStackReg (v1.46.5) plugin to apply a rigid-body transformation that aligned the Cy3 and Cy5 images to the corresponding DAPI image in a representative field of view. Transformations were applied across all fields of view in each experiment, and both the *E. coli* and sediment experiments used dual-labelled *E. coli* as the reference for extracting the transformation matrix. Next, for *E. coli*, we used the DAPI channel as our reference to find ROIs after applying the Moments automatic threshold, and we only selected those ROIs that had areas between 0.1-2 µm^2^. These ROIs were then redirected onto the corresponding raw DAPI, Cy3, and Cy5 images to extract the mean cell-specific intensity across each channel. For the salt marsh sediments, we chose to use the Cy3 and Cy5 channels (with a Moments threshold) as our reference to find ROIs, since the diffuse DAPI background in many of these images would lead to false positives/negatives with automated thresholding methods. We subsequently applied a Watershed (tolerance: 0.5) to separate overlapping cells and kept ROIs with areas between 0.1-4 µm^2^ and a circularity between 0.2-1.

Normalizing the cell-specific fluorescence in each channel by the corresponding DAPI signal is often useful to account for slight variations in intensity for cells on the edge of the focal plane. However, the cell-specific DAPI fluorescence varied dramatically in the salt marsh sediments (likely due to species-specific variations in GC content (36, 37); genome size, substrate limitation, growth stage, and number of genome copies (38); and permeability (39, 40)), leading to unreliable “normalized” intensities. (These factors also precluded an analysis of Cy3:Cy5 ratios for confirmed DAPI-active cells comparable to that conducted for *E. coli* cells, e.g., Fig. S7.) Therefore, we report the raw mean cell-specific intensities for all analyses, which remain comparable across experiments since the same imaging parameters were used throughout the study (Figs. 1, 3, S3, S10). To identify differentially labelled cells, we considered only those cells whose mean intensities in either the Cy3 or Cy5 channels were greater than the ncAA-free (but dyed) controls as being labelled with the corresponding ncAA. For a given cell, if both intensities were greater than the ncAA-free condition, then it was considered “dual-labelled.” Otherwise, it was either “only HPG” or “only AHA” labelled (Fig. S7).

All plots in this study were made using ggplot2 (v3.4.4 https://doi.org/10.32614/CRAN.package.ggplot2). Statistical significance was determined using the Whitney-Mann *U* test (R function *wilcox.test*). False positives rates in the dual-BONCAT validation experiments were calculated based on the portion of cells in the mono-labelled controls that were dimmer than the maximum mean cell fluorescence in either the ncAA-free or opposite mono-labelled controls, depending on which was greater. Additional packages used for data formatting and plotting include the following:

dplyr (v1.1.4 https://doi.org/10.32614/CRAN.package.dplyr)

reshape2 (v1.4.4 https://doi.org/10.32614/CRAN.package.reshape2)

stringr (v1.5.0 https://doi.org/10.32614/CRAN.package.stringr)

extrafont (v0.19 https://doi.org/10.32614/CRAN.package.extrafont)

scales (v1.2.1 https://doi.org/10.32614/CRAN.package.scales)

tidyr (v1.3.1 https://doi.org/10.32614/CRAN.package.tidyr)

rstudioapi (v0.17.1 https://doi.org/10.32614/CRAN.package.rstudioapi)

gg4hx (v0.3.1.9000 https://doi.org/10.32614/CRAN.package.ggh4x)

forcats (v1.0.1 https://doi.org/10.32614/CRAN.package.forcats)

### Fluorescence-Activated Cell Sorting

#### BONCAT Protocol Modifications for FACS

Because FACS requires stained cells to be in suspension, some modifications to the slide-based protocol described above were implemented. Starting from fixed cells in a 1:1 solution of PBS:ethanol, a 5-minute centrifugation at 16,000 x *g* was performed to pellet the cells, and the supernatant was discarded. The cells were resuspended in 250 µL of 80% ethanol, vortexed, and incubated at room temperature for 3 minutes. Next, 1.5 mL of 96% ethanol was added to the tube, which was then vortexed and left at room temperature for 3 minutes. The sample was centrifuged at 16,000 x *g* for 5 minutes, the supernatant was removed, and the pellet was resuspended in 221 µL of 1x PBS. The cells were pelleted again (16,000 x *g* for 5 minutes, supernatant removed) and resuspended in 500 µL of 100 mM 2-chloroacetamide in 1x PBS, and incubated for 1 hour in the dark. To stain the azide AHA molecules, DBCO-Cy5 was added to reach the desired final concentration (1 µM for cells extracted from salt marsh sediment to be used for FACS), and the sample was incubated in the dark at 46 °C for 30 minutes. Following the incubation, the sample was washed three times with PBS and once with 50% ethanol in 1x PBS; each wash involved centrifuging at 16,000 x *g* for 5 minutes, removing the supernatant, and resuspending cells in wash fluid. After the final wash step, the sample was resuspended in 500 µL of 1x PBS solution containing the desired concentration of the azide-Cy3 dye (25 µM, in this case). The tube was inverted once to gently mix, and the sample was left to incubate in the dark at room temperature for 30 minutes. To remove any unbound dye, the sample was then washed three times with 1x PBS and once with 50% ethanol in 1x PBS; each wash involved centrifuging at 16,000 x *g* for 5 minutes, removing the supernatant, and resuspending cells in wash fluid. Finally, the cells were resuspended in a 1:1 solution of PBS:ethanol and kept at –20 °C prior to loading onto the FACS instrument. For microscopic confirmation of staining, a few microliters can be placed on a slide and counterstained with DAPI, as described above.

#### Cell Sorting Procedures

Six samples of extracted salt marsh sediment microbial communities were sent to the University of Arizona for fluorescence-activated cell sorting (FACS): Condition 1 (ncAA#1: HPG-azide-Cy3; ncAA#2: AHA-DBCO-Cy5); Condition 2 (ncAA#1: AHA-DBCO-Cy5; ncAA#2: HPG-azide-Cy3); Cy3 Positive Control (mono-BONCAT with HPG-azide-Cy3); Cy5 Positive Control (mono-BONCAT with AHA-DBCO-Cy5); Cy3&Cy5 Positive Control (both ncAAs added for the full duration of an incubation experiment); and Negative Control (neither ncAA was added, but azide-Cy3 and DBCO-Cy5 dyes were). A BD Influx flow cytometer (BD Life Sciences, USA) was used to isolate cells from distinct gated subpopulations by FACS. The instrument fluidic system was thoroughly cleaned before use with multiple rinses of 10% bleach, 70% ethanol, and ultrapure water (8). The sheath fluid consisted of 1x PBS filtered through a 0.2-µm filter and autoclaved at 121°C for 30 min (hereafter referred to as sterile). The sheath fluid was also filtered through a 0.2-µm Sterivex filter (FisherScientific catalog #SVGPL10RC) connected in line with the sheath tank. A 70-μm nozzle was used, with a sheath fluid pressure of 30 PSI (207 kPa) and a sample fluid pressure of 31 PSI (214 kPa). The instrument was aligned, focused, and calibrated using reference microspheres (1-µm Fluoresbrite Multifluorescent, Polysciences catalog #24062; and 3-µm Ultra Rainbow, Spherotech catalog #URFP-30-2).

Samples were vortexed and filtered onto a BD Falcon 40-μm nylon cell strainer (BD Biosciences, catalog #352340) into polypropylene disposable tubes (BD Biosciences, catalog #352063) to avoid clogging the nozzle with cell clumps. To identify populations of interest, cell doublets were initially excluded using a trigger pulse width versus forward scatter plot. This approach assumes that pulse width typically covaries with forward scatter for single particles; doublets exhibit comparable forward scatter to single cells but have an increased pulse width. Consequently, total events were gated to include only those with similar pulse width to forward scatter ratios in the analyses (Fig. S13). A scatter plot displaying 585/29 nm emission at 561 nm excitation versus forward scatter was used to detect Cy3 labeled cells (hereafter referred to as the Cy3 plot), while a scatter plot featuring 692/40 nm emission at 640 nm excitation versus forward scatter detected Cy5 labeled cells (hereafter called the Cy5 plot). Another scatter plot was generated to detect Cy3&Cy5 dual-labeled cells (hereafter called the Cy3&Cy5 plot), showcasing 692/40 nm emission at 640 nm excitation (Cy5) versus 585/29 nm emission at 561 nm excitation (Cy3). Negative controls consisting of ncAA-free samples with Cy3, Cy5, or Cy3 and Cy5 dyes were used to draw the background gates on the Cy3, Cy5, and Cy3&Cy5 plots. Cy3 and Cy5 mono-labeled control samples were analyzed to determine the background signal generated in the Cy5 and Cy3 plots, respectively (i.e., to determine fluorescence bleed-through), and to draw the Cy3-positive and Cy5-positive subpopulation gates. Cy3&Cy5 dual-labeled control samples were analyzed to draw the Cy3&Cy5-positive gate on the Cy3&Cy5 plot.

At the beginning of each sample processing day, the drop delay was calibrated using BD FACS Accudrop Beads (catalog #345249). The sorting efficiency was manually verified by sorting a designated number of 1-μm Fluoresbrite yellow-green microspheres (Polysciences catalog #17154-10) onto a glass slide, followed by counting the beads under an epifluorescence microscope. The breakoff position was monitored throughout the day and adjusted as necessary. The instrument was set to high sorting purity (1.0 drop) mode. To maintain high sorting purity and recovery, samples were diluted in sterile 1x PBS as needed to achieve an event rate of less than 5,000 per second, thereby avoiding coincidence and swarm (41). Cells from each sorted subpopulation were collected into separate sterile microtubes. After collection, samples were stored frozen at –80°C. The number of sorted cells recovered using the approaches described above is provided in Table S1.

### 16S rRNA Gene Sequencing & Analysis

#### DNA Extraction & Sequencing

DNA was extracted from the FACS-sorted samples by a modified hot phenol/chloroform extraction (42, 43). All reagents were molecular grade and nuclease-free. Sorted samples were added to Invitrogen Bead Tubes (Thermo Fisher #A29158) with 500 µL of extraction buffer (10% (w/v) CTAB buffer at pH 8, 100 mM Tris-HCl, 100 mM Na-EDTA, 100 mM NaH_2_PO_4_, 1.5 M NaCl) heated to 75°C (for easier transfers). Bead tubes were vortexed for 5 minutes and inverted periodically to ensure proper mixing. 50 µL of 50 mg/mL lysozyme and 10 µL of 20 mg/mL Proteinase K were added to each tube and incubated at 37 °C for 30 minutes. After incubation, 210 µL of 20% (w/v) sodium dodecyl sulfate (SDS) were added to each bead tube and incubated for 2 hours at 65 °C. Tubes were then centrifuged at 10,000 x *g* for 10 minutes and the supernatant was transferred to a 2 mL Eppendorf tube and placed on ice. A second extraction was performed by adding an additional 500 µL of extraction buffer to the original bead tubes. Bead tubes were vortexed for another 10 minutes and inverted periodically. Another 70 µL 20% (w/v) SDS was added and tubes were incubated at 65 °C for 1 hour. Bead tubes were centrifuged at 10,000 x *g* for 10 minutes, and the remaining supernatants were combined with the first extractions. An equal volume of 60 °C (pre-heated) phenol:chloroform:isoamyl alcohol (25:24:1, adjusted to pH 7.9 with Tris buffer) was added to each extract and inverted gently to mix. Tubes were centrifuged at 13,800 x *g* for 30 minutes, and the upper aqueous layer was transferred into a fresh 1.5 mL Eppendorf tube. To precipitate the DNA, approximately 0.6 extract volume’s worth of isopropyl alcohol and 0.6 volume’s worth of 0.3 M Na-Acetate (pH 5.2) were added to each Eppendorf tube and inverted gently to mix. Tubes were placed at 4 °C and allowed to precipitate overnight. Tubes were centrifuged at 16,000 x *g* for 20 minutes, and the supernatant was discarded. DNA pellets were washed with ∼200-500 µL of chilled 75% ethanol and centrifuged for another 16,000 x *g* for 10 minutes. The supernatant was discarded and the remaining DNA pellets were allowed to air dry for 15-30 minutes. DNA was resuspended in 50 µL of nuclease-free water and stored at –20 °C. DNA was quantified on a Qubit 4 Fluorometer (Invitrogen) following the manufacturer’s instructions.

Extracted DNA was sent to the University of Delaware DNA Sequencing and Genotyping Center for library preparation and sequencing. Initially, some samples did not generate a library, so all samples were cleaned using a Zymo Clean & Concentrator-5 kit. The V4 region of the 16S rRNA gene was amplified based on the Earth Microbiome protocol (44) with the revised 515 F (5’-GTGYCAGCMGCCGCGGTAA-3’; (45)) and 806 R (5’-GGACTACNVGGGTWTCTAAT-3’; (46)) primers for 30 cycles and sequenced on an Illumina MiSeq 251-cycle paired-end run (2×251) with an output of 2M paired reads (analyses only performed on the forward reads, see below). Two controls were included in the sequencing run: a “DNA Extraction Negative Control” that followed the extraction procedure detailed above but included no template, and a “UDel Sequencing Negative Control” that resulted from running 31 µL H_2_O through the clean and concentrate column and using it as the PCR negative control.

#### Sequence Analysis

Raw sequence reads were processed on the Boston University Shared Computing Cluster using the BU16s pipeline (https://github.com/Boston-University-Microbiome-Initiative/BU16s) built with QIIME 2020.2 (47) as described in (48) and (49). Sequencing adaptors were removed with cutadapt (50) and amplicon sequence variants (ASVs) were generated with dada2 (51). For taxonomic association, ASVs were clustered to 99% identity with the SILVA 132 database (52) using the vsearch *cluster-features-closed-reference* (53). Initially, reads were analyzed as paired-ends, but this led to few reads passing through the quality filter. Poor reverse read quality was confirmed with FastQC, and the forward reads were processed as a single-end library for subsequent analyses. Raw sequence reads have been archived at the National Centre for Biotechnology Information GenBank under the Bioproject ID PRJNA1188625.

ASV analyses were performed with R (v4.3.1) in RStudio (v2023.09.1+494). Alpha diversity metrics were calculated as described in (11) and are provided in Table S8. Data was formatted using Phyloseq (v1.46.0; (54)); to compare samples with uneven sequencing depth, ASVs were rarefied with GUniFrac (v1.8 https://doi.org/10.32614/CRAN.package.GUniFrac). Species richness (Chao and Abundance-based Coverage Estimator (ACE) metrics) and the Shannon diversity index were calculated by Vegan using the *estimateR* and *diversity* functions, respectively (v2.6-10 https://doi.org/10.32614/CRAN.package.vegan). Evenness was determined by dividing the Shannon diversity index by the natural-logarithm of the number of observed ASVs.

Relative abundances were calculated by dividing the number of reads from each ASV by the total number of reads in each sample (Table S3). Ambiguous taxonomic classifications (e.g. uncultured, unidentified etc.) were removed from each ASV up to the lowest defined category. For family-level analyses, ASVs were grouped at the family level (or their lowest defined taxonomic label) and relative abundances within each family-level grouping were summed together. If the family was not known, the remaining ASVs were grouped by their order, and so on, moving further up the taxonomic ladder until the domainless ASVs were grouped (Table S2). Of this data, we considered the grouped taxa which accounted for >1% relative abundance in at least one of the conditions (Fig. 5; Table S2). The mean and standard deviation of the Day-active (ncAA#1) and Day– and Night-active or Night-active (ncAA#1 & #2) families were calculated from the two experimental conditions and plotted alongside the Night-active (ncAA#2) families from Condition 2 using ggplot2 (v3.4.4 https://doi.org/10.32614/CRAN.package.ggplot2) (Fig. 5). Additional packages used for data formatting and plotting include the following:

dplyr (v1.1.3 https://doi.org/10.32614/CRAN.package.dplyr)

reshape2 (v1.4.4 https://doi.org/10.32614/CRAN.package.reshape2)

stringr (v1.5.0 https://doi.org/10.32614/CRAN.package.stringr)

extrafont (v0.19 https://doi.org/10.32614/CRAN.package.extrafont)

scales (v1.2.1 https://doi.org/10.32614/CRAN.package.scales)

### Data Availability

All datasets generated during and/or analyzed during the current study are available from the corresponding author upon reasonable request. Raw sequence reads have been archived at the National Centre for Biotechnology Information GenBank under the Bioproject ID PRJNA1188625.

## Acknowledgments

We would like to thank Paul Bae and Theodore Dowd for assisting with lab procedures; Claire Andrade, Erin Frates, Jiayi Li, Vinitra Nathan, Dr. Paul Rousteau, and Peter Schroedl for reviewing data products; Dr. Roland Hatzenpichler for reviewing an earlier version of the manuscript; Brewster Kingham and Mark Shaw at the University of Delaware DNA Sequencing and Genotyping Center for sequencing services and troubleshooting; and Thomas Symancyk and Zachary Jones for logistical and shipment support. This work was performed under the auspices U.S. Department of Energy Grant #DE-SC0022193 and NASA Exobiology Grant #80NSSC23K0224.

---

**Fig. S1:**
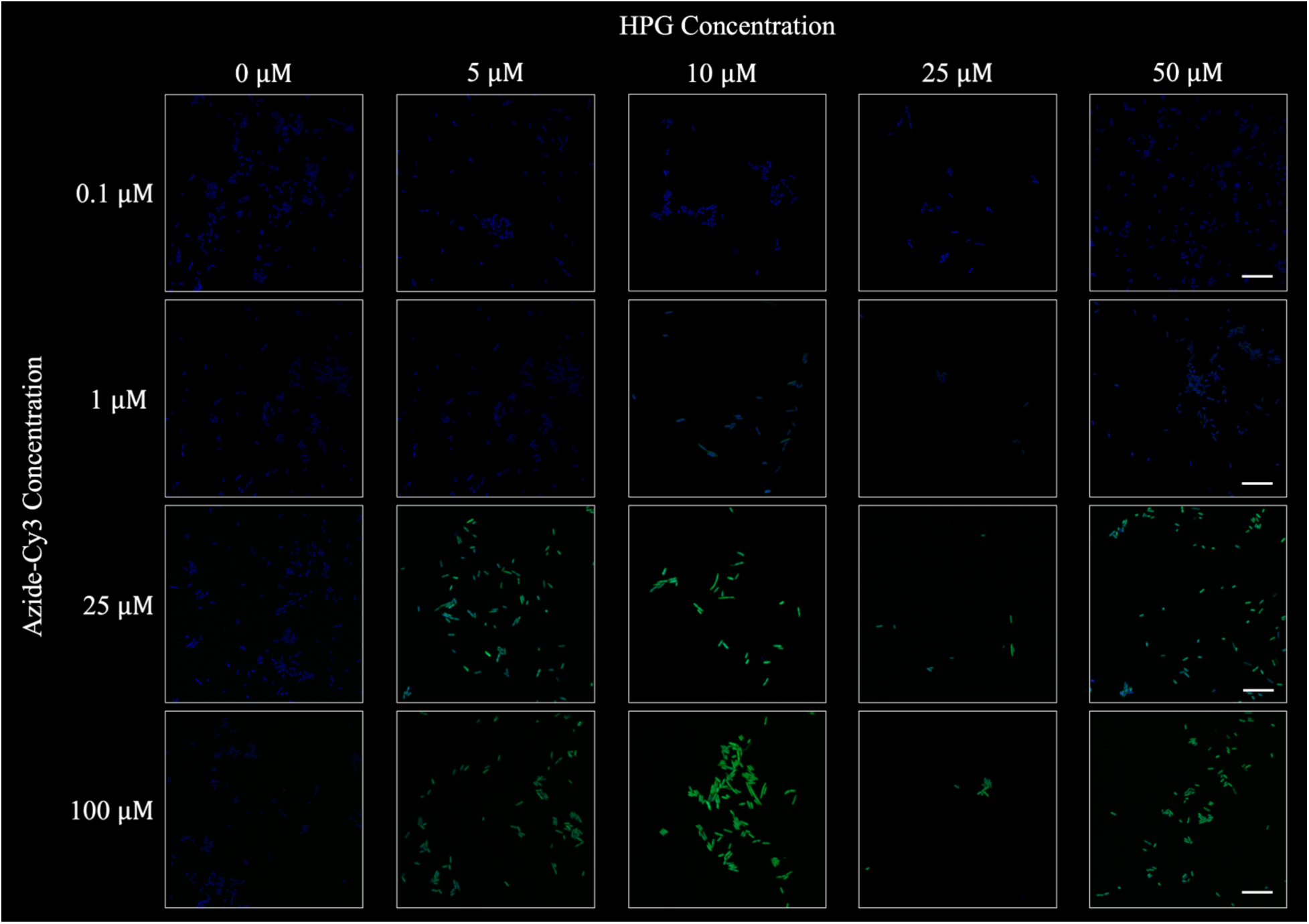
Representative fields of view from mono-BONCAT experiments with *E. coli* cultures, evaluating fluorescence signals derived from a range of HPG and Azide-Cy3 concentrations. All scale bars are 10 µm.

**Fig. S2:**
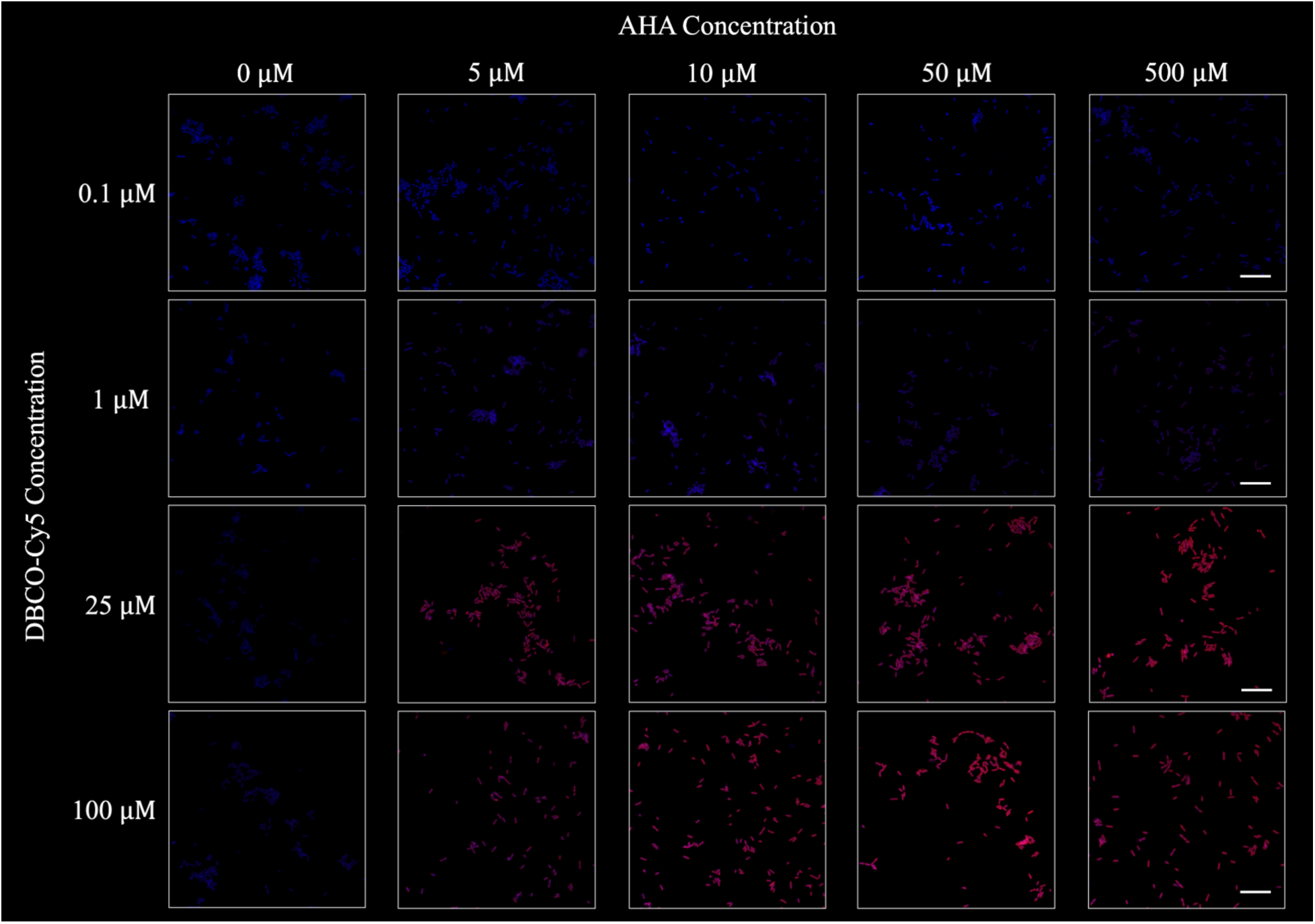
Representative fields of view from mono-BONCAT experiments with *E. coli* cultures, evaluating fluorescence signals derived from a range of AHA and DBCO-Cy5 concentrations. All scale bars are 10 µm.

**Fig. S3:**
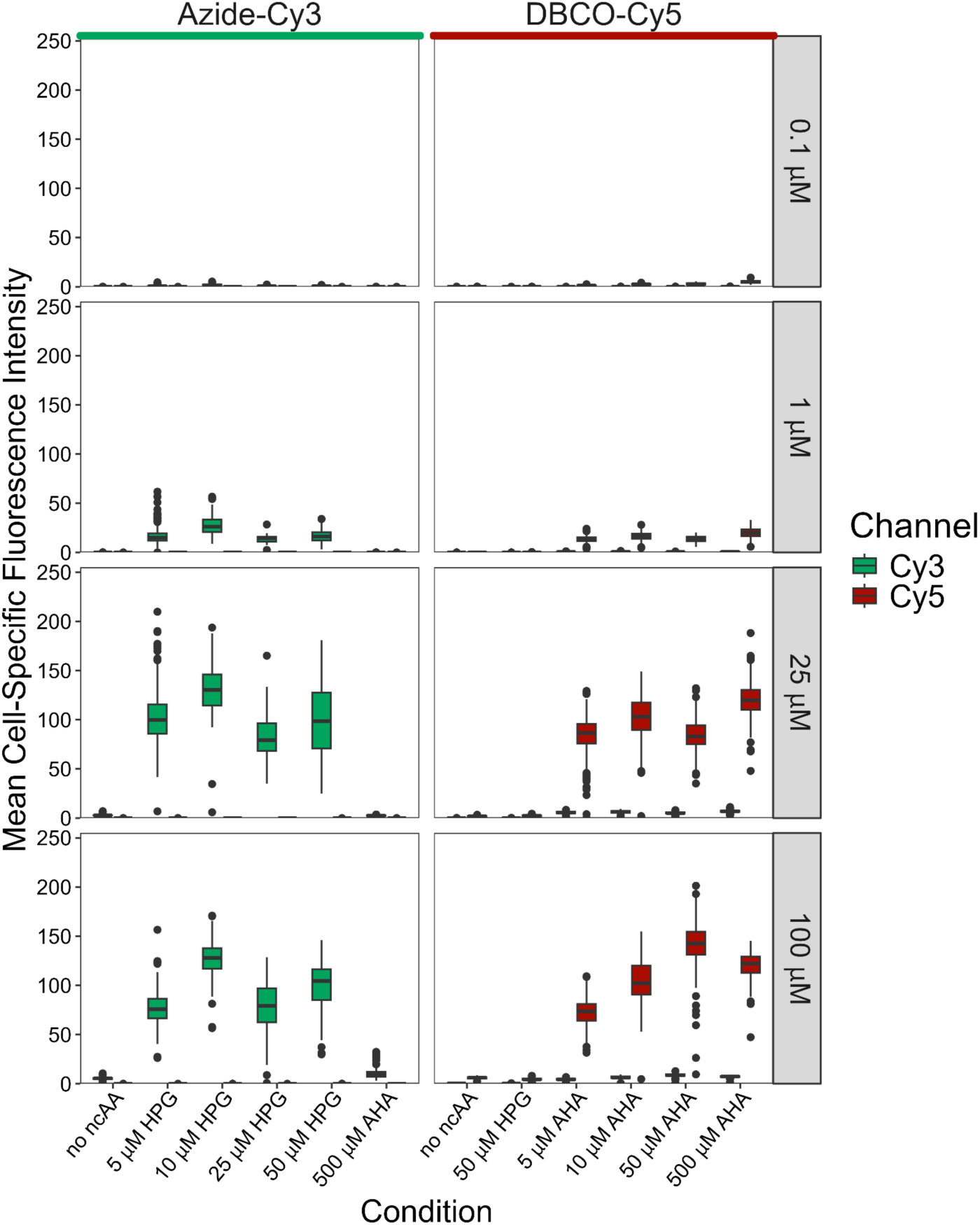
Boxplots of mean fluorescence intensity values of cells in mono-BONCAT experiments with *E. coli* cultures. A range of ncAA and dye concentrations were tested to determine parameters for subsequent dual-BONCAT experiments. Imaging parameters were consistent across all different conditions.

**Fig. S4:**
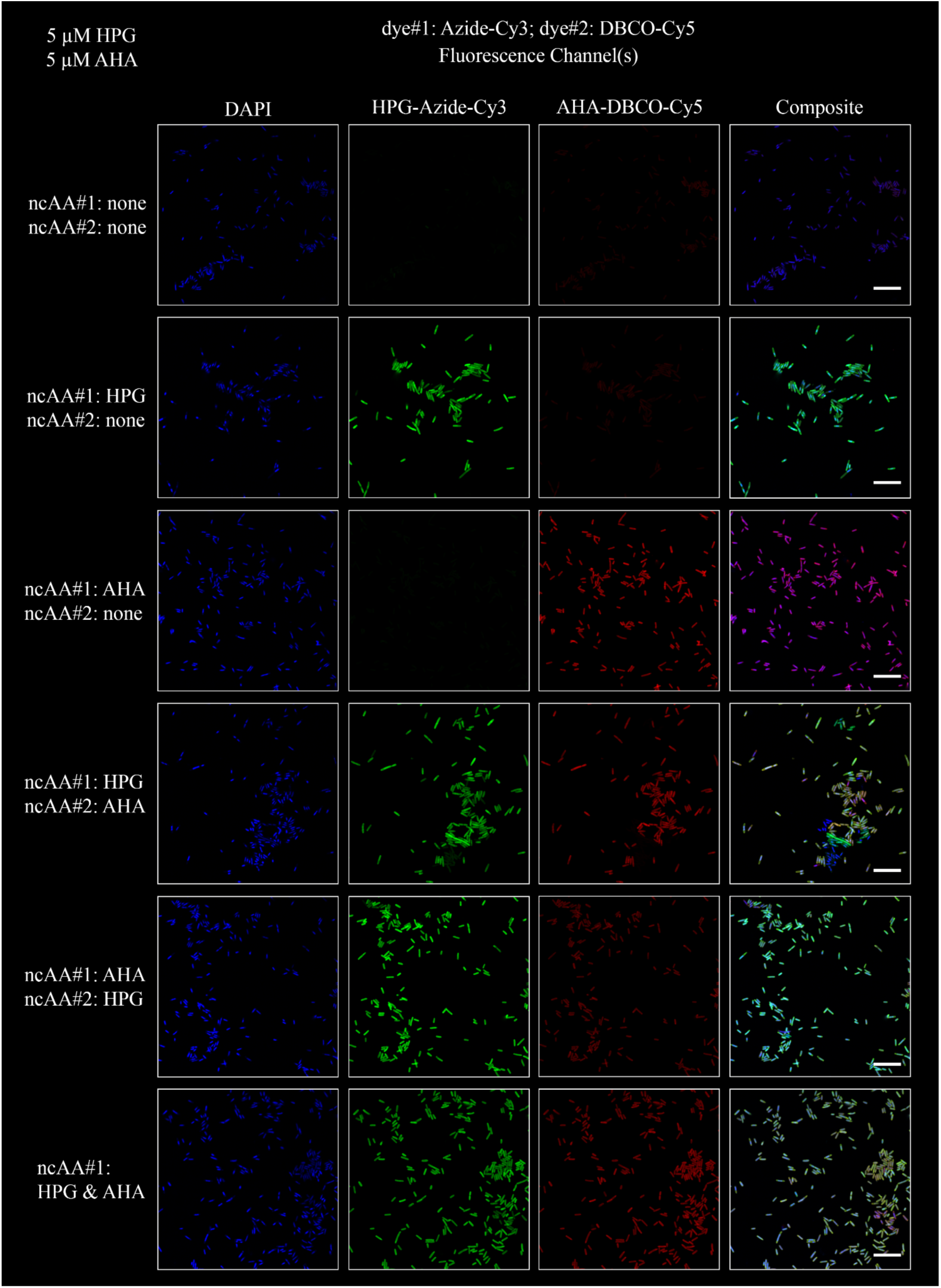
Representative fields of view from dual-BONCAT experiments with *E. coli* cultures, evaluating the order of ncAA addition and performing the HPG-Azide reaction first followed by the AHA-DBCO. All scale bars indicate 10 µm.

**Fig. S5:**
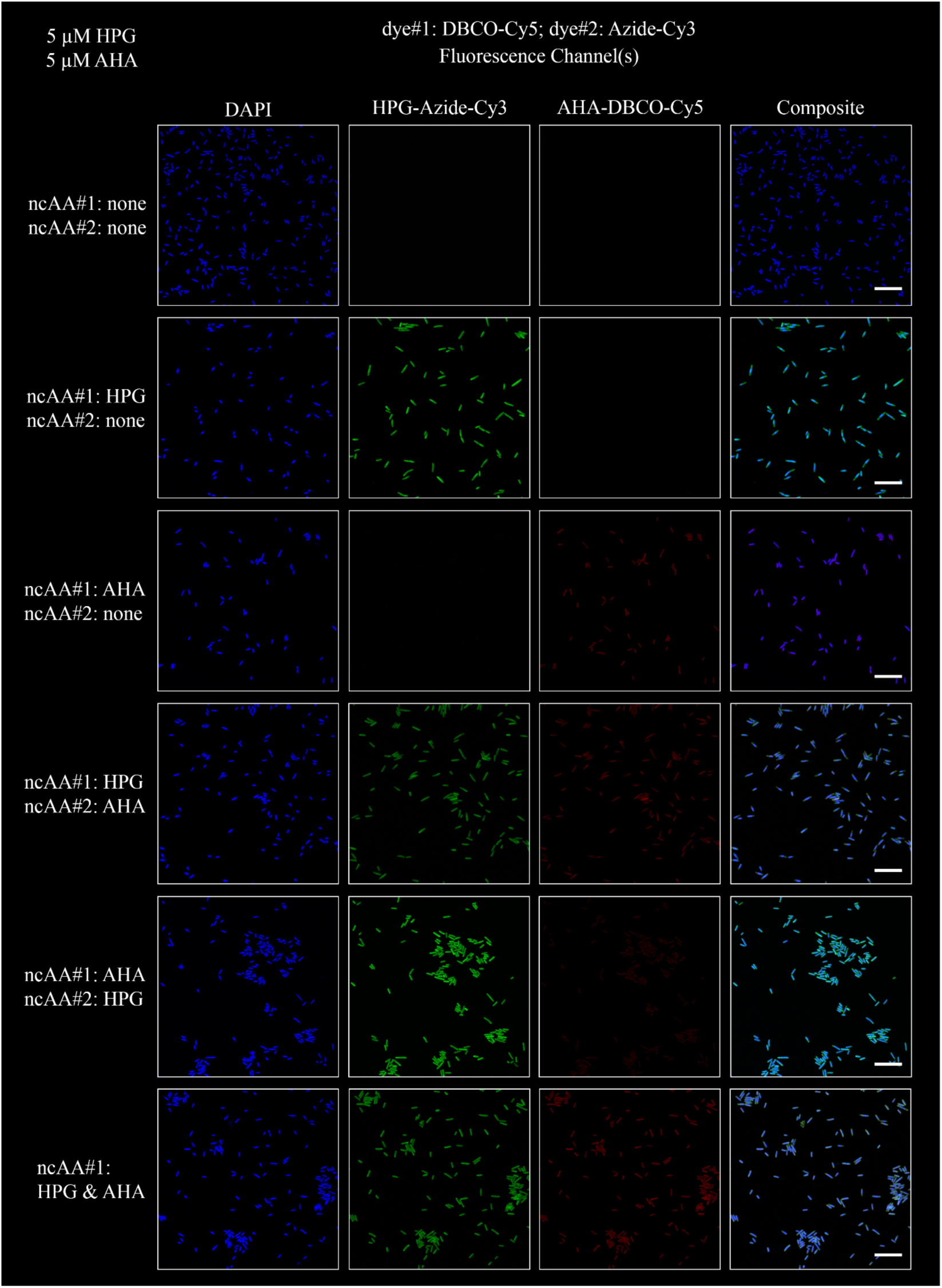
Representative fields of view from dual-BONCAT experiments with *E. coli* cultures, evaluating the order of ncAA addition and performing the AHA-DBCO reaction first followed by the HPG-Azide reaction. All scale bars indicate 10 µm.

**Fig. S6:**
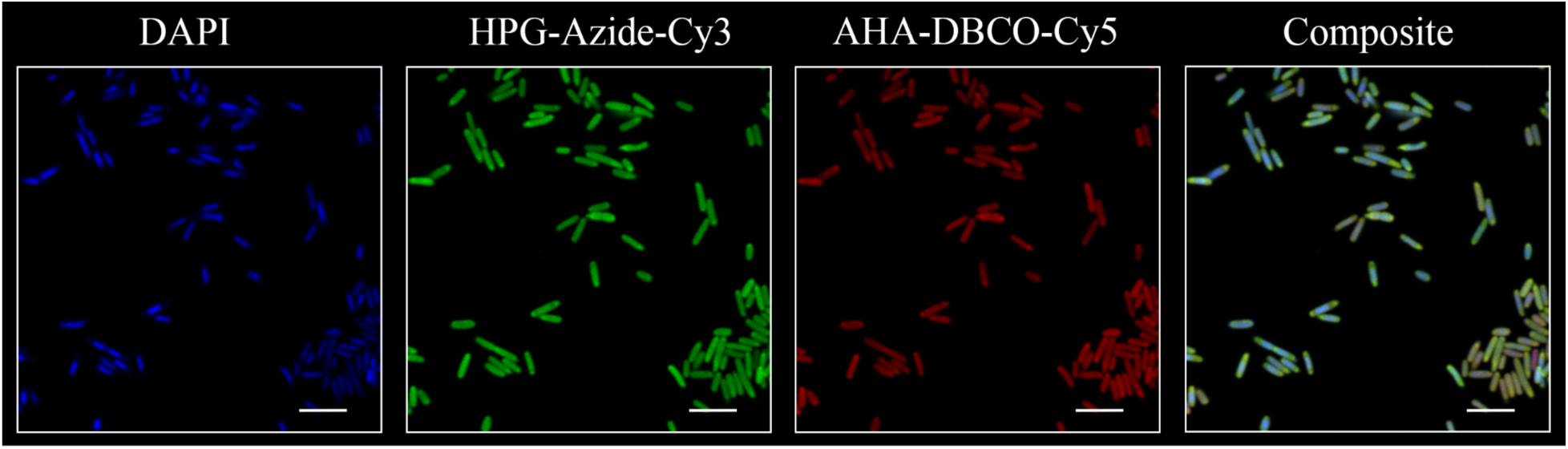
Fluorescence microscopy field of view of a dual-BONCAT experiment with *E. coli* cells. Both ncAAs were added at the start of the experiment, and, following fixation, the Azide-Cy3 click reaction was performed first. Scale bar indicates 5 µm.

**Fig. S7:**
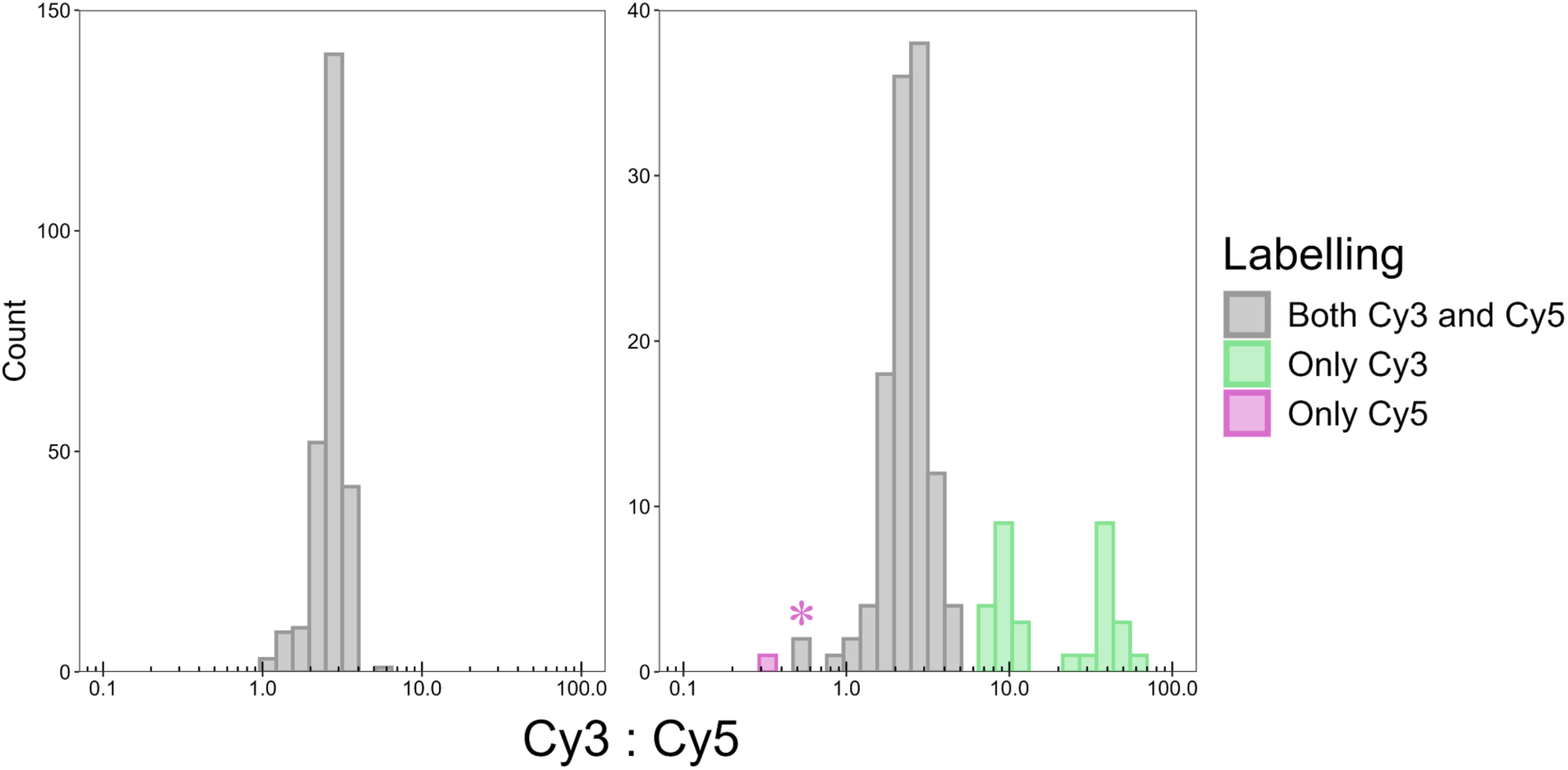
Ratio of the Cy3 to Cy5 fluorescent intensities for the experiment in which HPG and AHA were added at T_1_ in the field of view chosen for Fig. S6 (left), and the experiment in which HPG was added at T_1_ and AHA at T_2_ in the field of view chosen for Fig. 2 (right). Cells whose mean intensities exceeded the maximum intensities in the “no ncAA” condition were considered labelled. The red-colored bar indicates the cell labeled with a red arrow in Fig. 2; the bar marked with a red asterisk indicates the two cells labeled with red arrows and asterisks in Fig. 2.

**Fig. S8:**
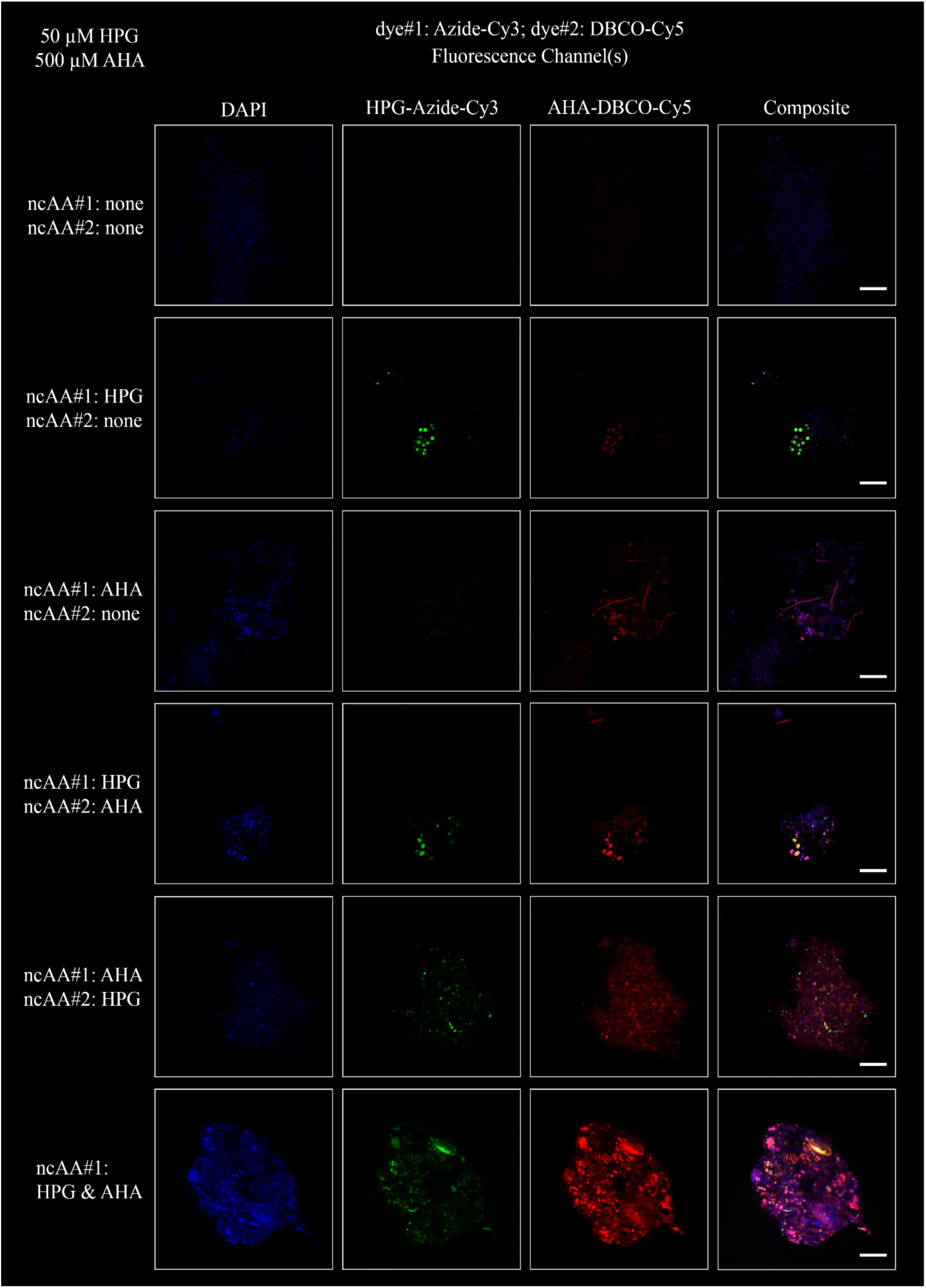
Representative fields of view from dual-BONCAT experiments with extracted cells from salt marsh sediments, evaluating the order of ncAA addition and performing the HPG-azide reaction first followed by the AHA-DBCO. All scale bars indicate 10 µm.

**Fig. S9:**
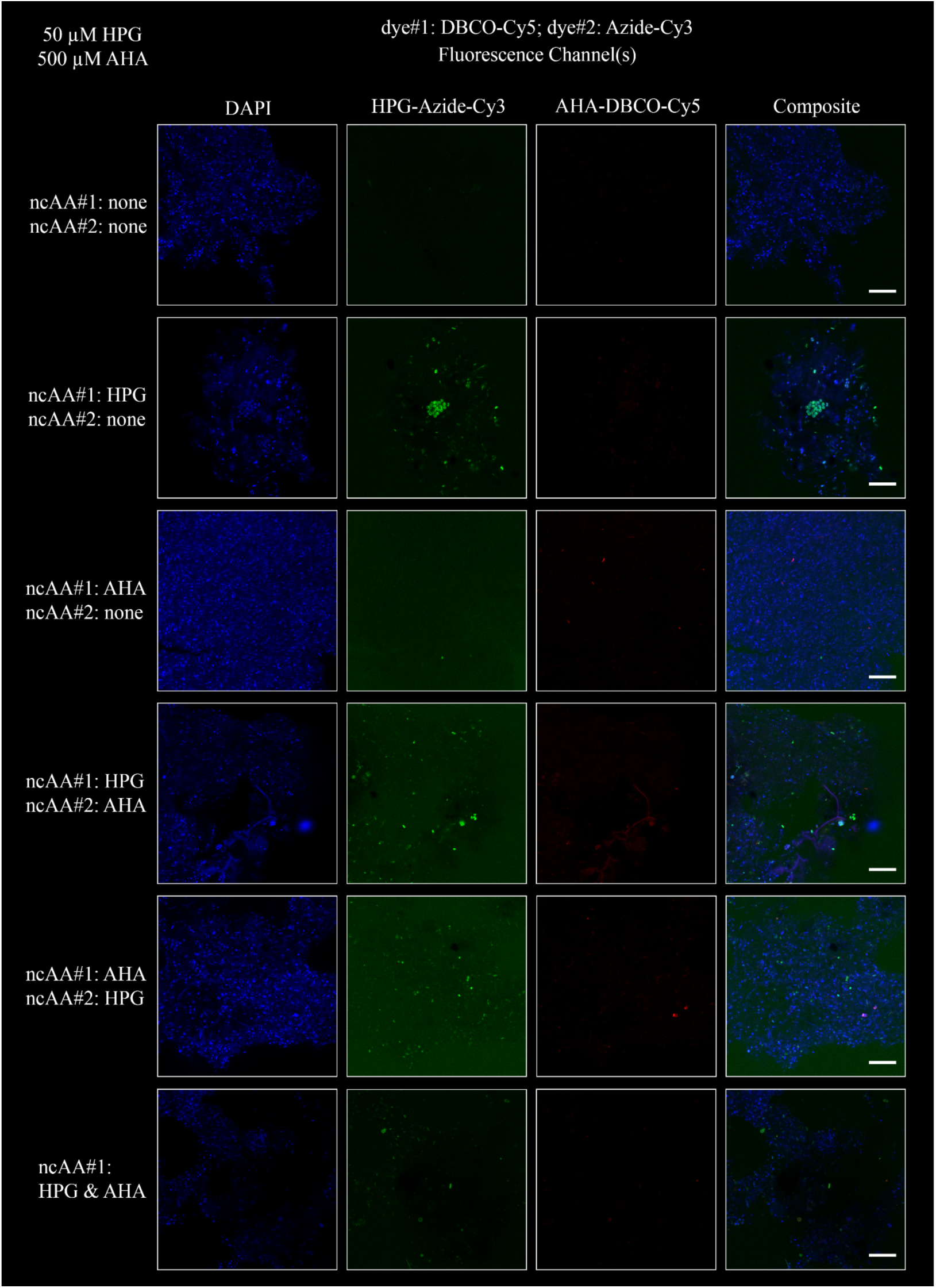
Representative fields of view from dual-BONCAT experiments with extracted cells from salt marsh sediments, evaluating the order of ncAA addition and performing the DBCO-Cy5 reaction first followed by the HPG-azide. All scale bars indicate 10 µm.

**Fig. S10:**
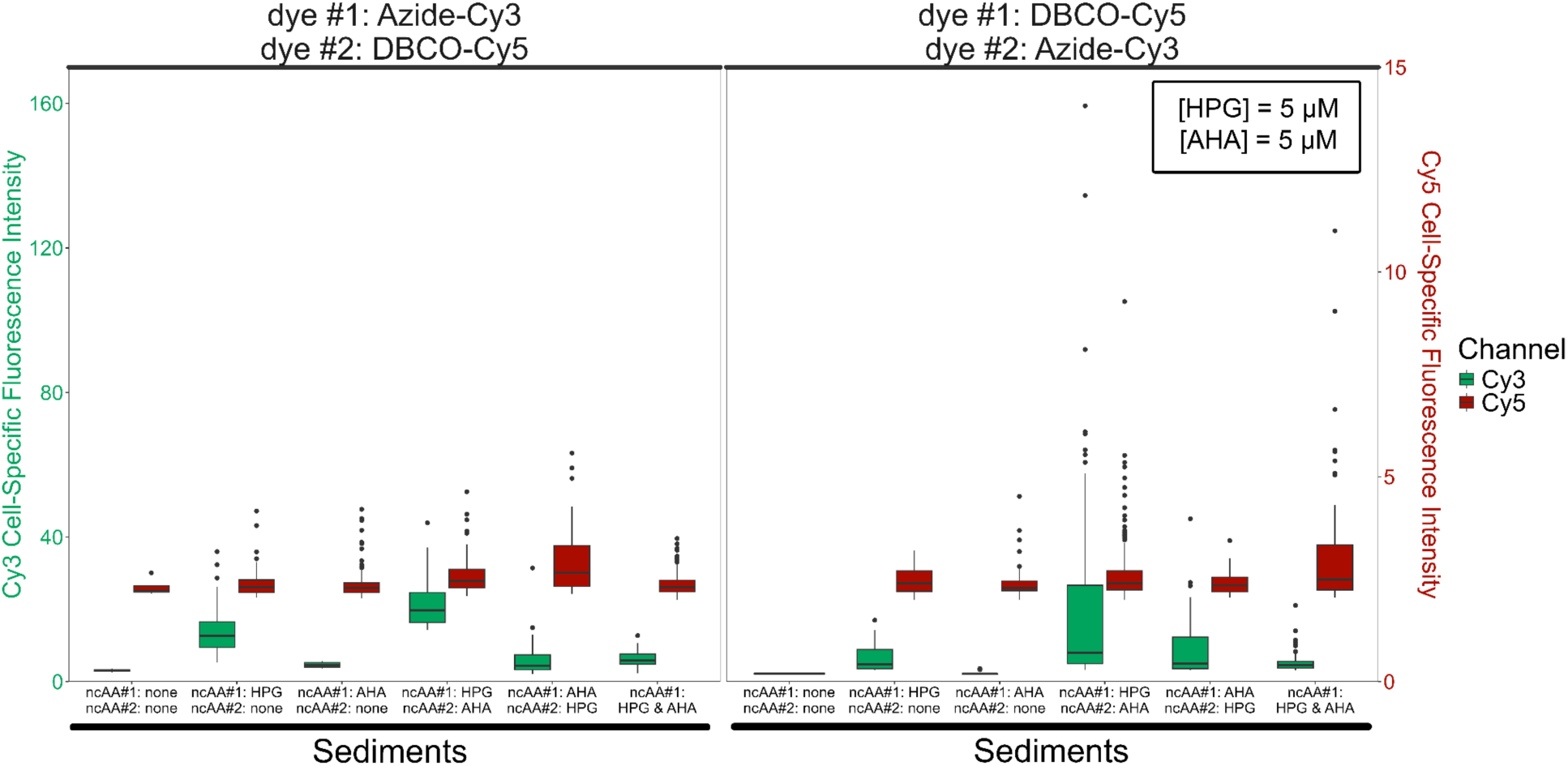
Boxplots of mean fluorescence intensity values of cells in dual-BONCAT experiments with extracted salt marsh sediment cells. Tests varying the sequence of ncAA addition (see column labels along x-axis) were performed, with the Azide-Cy3 click reaction conducted first (left panel) and with the DBCO-Cy5 click reaction conducted first (middle panel). Data from the 5 µM HPG & 5 µM AHA condition.

**Fig. S11:**
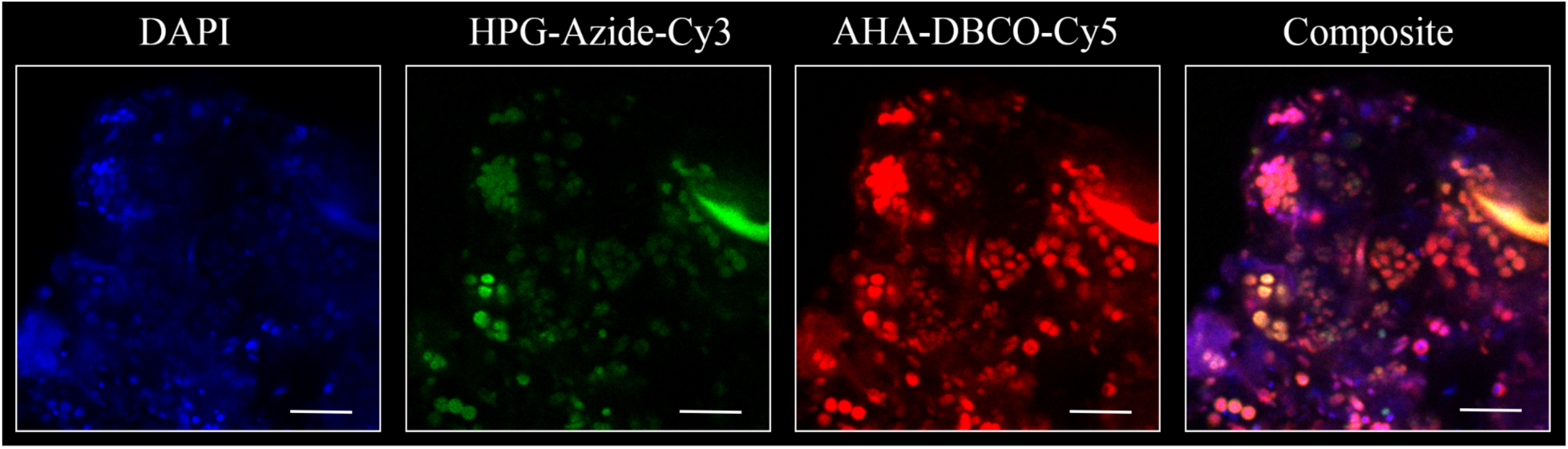
Fluorescence microscopy field of view of a dual-BONCAT experiment with extracted salt marsh sediment cells. Both ncAAs (50 µM HPG & 500 µM AHA) were added at the start of the experiment, which was stopped after a single, 24-hour day-night cycle. The Azide-Cy3 click reaction was performed first. Scale bar indicates 5 µm.

**Fig. S12:**
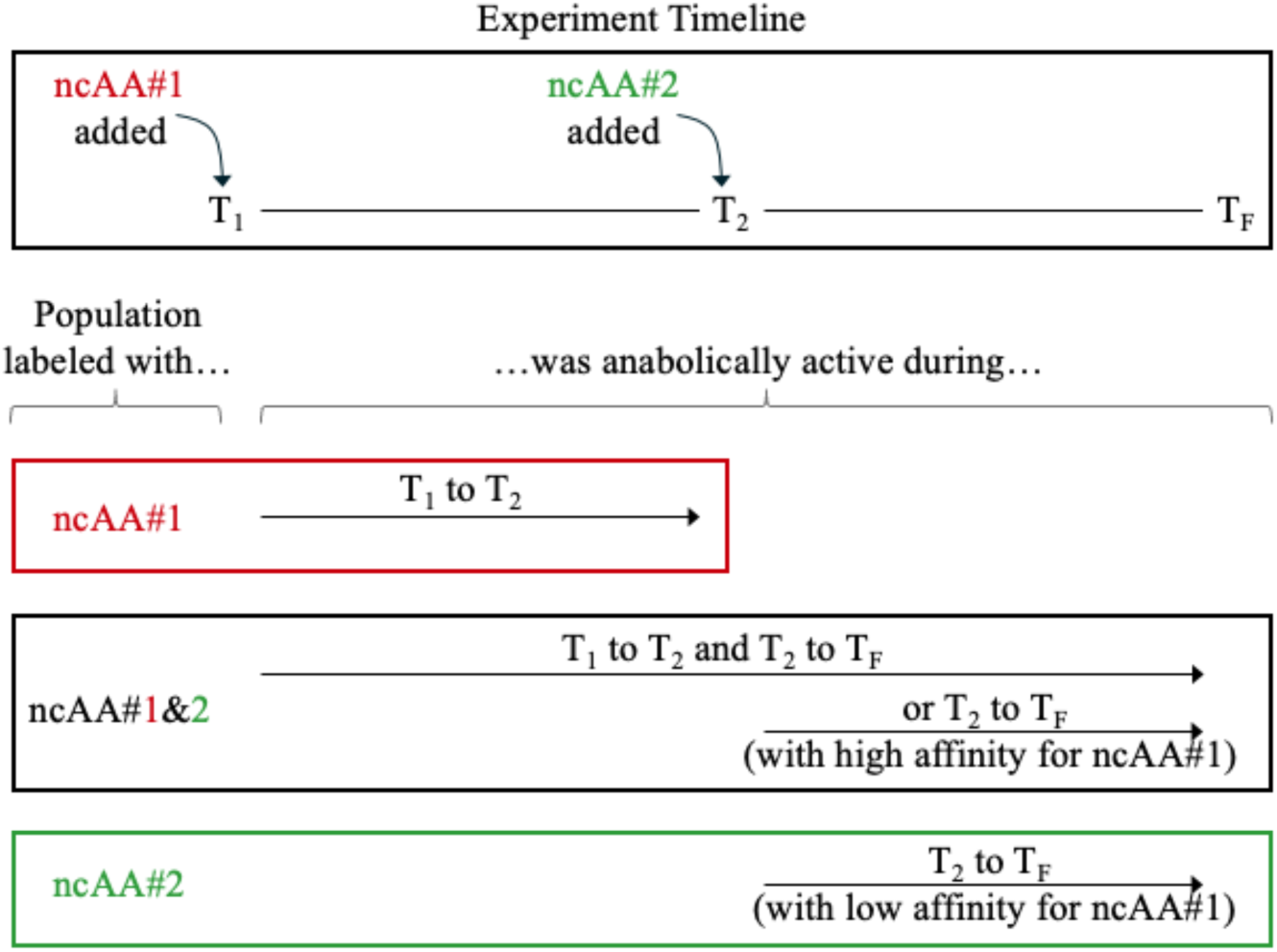
An interpretive scheme for dual-BONCAT experiments. For a generalized experimental timeline during which one ncAA is added at an initial time point (T_1_) and a second ncAA is added at a subsequent time point (T_2_), the three distinct subpopulations of labeled cells (e.g., those labeled with only ncAA#1, both ncAA#1&2, and only ncAA#2) can be interpreted as being anabolically active during distinct phases of the experiment. Further details are provided in the text.

**Fig. S13:**
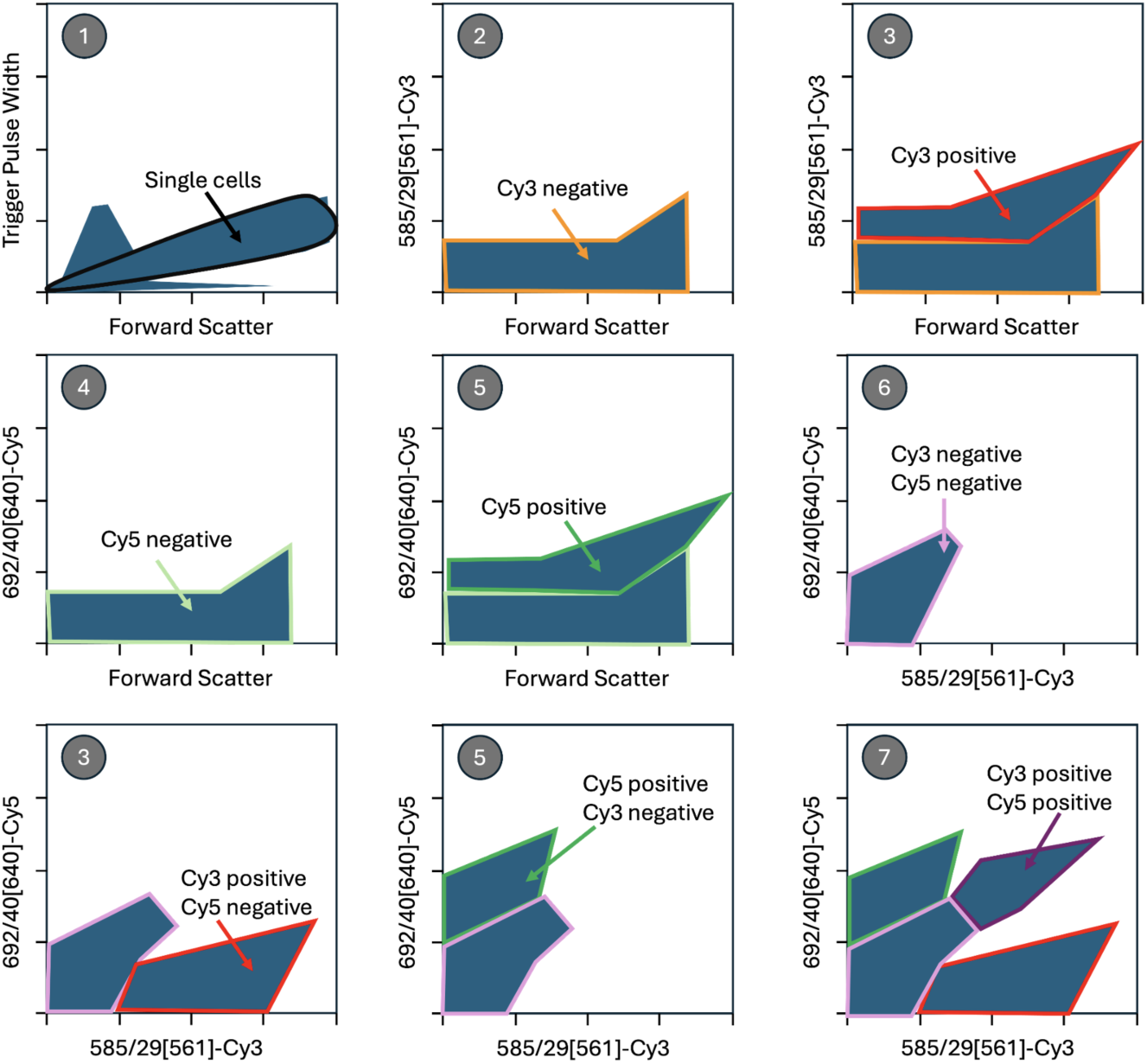
Schematic representation of the subpopulations gating protocol. (1) Gate single cells subpopulation on a trigger pulse width versus forward scatter plot, (2) Gate Cy3 negative subpopulation by analyzing a ncAA-free sample with Cy3 and a Cy5 mono-labeled sample on the Cy3 plot [585/29 nm emission at 561 nm excitation versus forward scatter], (3) Gate Cy3 positive subpopulation by analyzing a ncAA-Cy3 sample on the Cy3 plot and the Cy3Cy5 plot [692/40 nm emission at 640 nm excitation versus 585/29 nm emission at 561 nm excitation], (4) Gate Cy5 negative subpopulation by analyzing a ncAA-free sample with Cy5 and a Cy3 mono-labeled sample on the Cy5 plot [692/40 nm emission at 640 nm excitation versus forward scatter], (5) Gate Cy5 positive subpopulation by analyzing a ncAA-Cy5 sample on the Cy5 plot and the Cy3Cy5 plot, (6) Gate Cy3&Cy5 double-negative subpopulation by analyzing a ncAA-free sample with Cy3 and Cy5 on the Cy3Cy5 plot, (7) Gate Cy3&Cy5 double-positive subpopulation by analyzing a ncAA-Cy3&Cy5 sample on the Cy3Cy5 plot.

**Table S1:** Number of sorted cells recovered from each gate of interest in this study.

| **Condition 1** |  |
| --- | --- |
| *Subpopulation* | *Number of Sorted Cells* |
| ncAA#1 active | 718,584 |
| ncAA#1&2 active | 295,495 |
| ncAA#2 active | Below Detection |
| **Condition 2** |  |
| *Subpopulation* | *Number of Sorted Cells* |
| ncAA#1 active | 21,424 |
| ncAA#1&2 active | 552,138 |
| ncAA#2 active | 321,110 |

**Table S2:** Taxonomic identity and relative abundance of ASVs grouped at the family level for the seven communities sequenced in this study. Only family-level groupings that accounted for >1% of the relative abundance of at least one sequenced community are shown. *See Excel File*

**Table S3:** Taxonomic identity and relative abundance of all ASVs for the seven communities sequenced in this study. *See Excel File*

**Table S4:** Components of the two premixes needed for the Cu(I)-catalyzed HPG-azide reaction.

| Premix 1 |  | Premix 2 |  |
| --- | --- | --- | --- |
| *Component* | *Volume* | *Component* | *Volume* |
| 100 mM Na-ascorbate | 20 μL | 20 mM CuSO<sub>4</sub> | 2 μL |
| 100 mM Aminoguanidine-HCl | 20 μL | 50 mM THPTA | 4 μL |
| 1x PBS | 344 μL | 1 mM azide dye | 10 μL |

**Table S5:** Imaging parameters for the culture and environmental samples used in this study.

| For *E. coli* cultures - 63x objective lens and 2.5x zoom |  |  |  |  |
| --- | --- | --- | --- | --- |
| Dye | Laser Excitation | Detector Emission Window | Laser Intensity | Detector Gain |
| DAPI | 405 nm | 420-559 nm | 7.0% | 8.0% |
| Cy3 | 554 nm | 571-704 nm | 2.0% | 2.5% |
| Cy5 | 649 nm | 666-750 nm | 1.5% | 2.5% |
| For LSSM Sediments - 63x objective lens and 2.5x zoom |  |  |  |  |
| Dye | Laser Excitation | Detector Emission Window | Laser Intensity | Detector Gain |
| DAPI | 405 nm | 420-559 nm | 7.0% | 2.5% |
| Cy3 | 554 nm | 571-704 nm | 2.0% | 2.5% |
| Cy5 | 649 nm | 666-750 nm | 1.5% | 2.5% |

**Table S6:** FIJI Multi-Channel Cell Counter plugin settings for the culture and environmental samples used in this study.

| For *E. coli* cultures |  |  |
| --- | --- | --- |
| Dye | Color | Min/Max |
| DAPI | Blue | [0,100] |
| Cy3 | Green | [0,200] |
| Cy5 | Red | [0,100] |
| Manually Adjusted? | Yes |  |
| Transformed? | Yes |  |
| For LSSM Sediments |  |  |
| Dye | Color | Min/Max |
| DAPI | Blue | [0,60] |
| Cy3 | Green | [0,30] |
| Cy5 | Red | [0,40] |
| Manually Adjusted? | Yes |  |
| Transformed? | Yes |  |

**Table S7:**
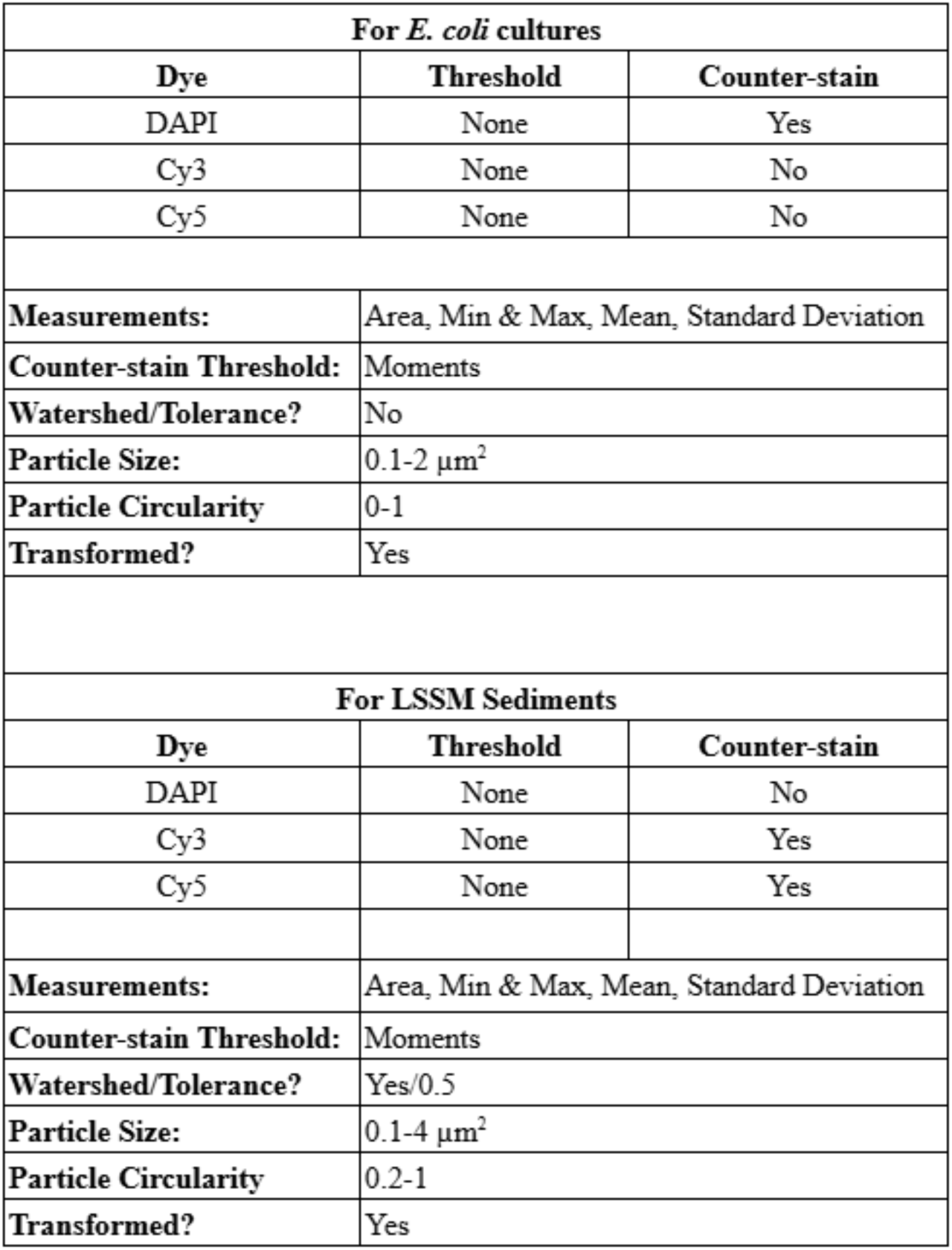
FIJI Multi-Channel RGB Stacker plugin settings for the culture and environmental samples used in this study.

**Table S8:** Alpha diversity metrics for the seven sequenced communities analyzed in this study.

| Sample | # of ASVs | Chao | ACE | Shannon | Evenness |
| --- | --- | --- | --- | --- | --- |
| Condition 1 Day-Active (ncAA#1) | 4212 | 4629.2 | 4588.3 | 6.74 | 0.81 |
| Condition 2 Day-Active (ncAA#1) | 687 | 707.8 | 704.1 | 4.23 | 0.65 |
| Condition 1 Day- & Night-active (ncAA#1&2) | 3256 | 3857.1 | 3811.6 | 5.98 | 0.74 |
| Condition 2 Day- & Night-active (ncAA#1&2) | 1891 | 2051.8 | 2060 | 4.70 | 0.62 |
| Condition 2 Night-active (ncAA#2) | 2352 | 2543.3 | 2479.5 | 5.56 | 0.72 |
| Delaware Negative Control | 22 | 22 | 22 | 1.32 | 0.43 |
| DNA Extraction Negative Control | 172 | 179 | 176.4 | 3.19 | 0.62 |

